# Tobacco Smoke Exposure is Characterized by a Distinct Nasal Rhinotype in a Pediatric Population

**DOI:** 10.1101/2024.03.15.585112

**Authors:** Hilary Monaco, Cordelia R Elaiho, Bian Liu, Tiffany Chan, Adam Cantor, Joseph Michael Collaco, Sharon A McGrath-Morrow, Karen M Wilson, Jose C Clemente

**Affiliations:** Department of Genetics and Genomics, Icahn School of Medicine at Mount Sinai; Department of Population Health Science and Policy, Icahn School of Medicine at Mount Sinai; Eudowood Division of Pediatric Respiratory Sciences, Johns Hopkins University School of Medicine; Division of Pulmonary Medicine, Children’s Hospital of Philadelphia and University of Pennsylvania; Department of Pediatrics, University of Rochester Medical Center; Medical College of Wisconsin; Department of Medicine, Division of Allergy and Immunology, Northwestern University

## Abstract

**Background:** Secondhand tobacco smoke exposure (TSE) increases susceptibility to respiratory diseases, but the mechanisms of action are poorly understood.

**Objective:** To study the effect of TSE in the nasal microbiome of children, and to evaluate whether such effect is dose-dependent with measured levels of cotinine in saliva and urine.

**Methods:** The study was performed at the Mount Sinai Kravis Children’s Hospital (New York, NY) and the Johns Hopkins Hospital (Baltimore, MD). We enrolled 236 children between 6 months and 10 years of age, both inpatients and outpatients. We collected swabs to characterize the diversity and composition of the nasal microbiome using 16S rRNA gene sequencing and measured cotinine levels in salivary and urinary samples to quantify TSE. We then determined the relationship between these measures and participant respiratory conditions, demographics and lifestyle factors.

**Results:** Infants with high cotinine levels had lower nasal microbiome alpha diversity and an enrichment in Moraxella, Dolosigranulum and Corynebacterium, which formed a distinct cluster in network analysis. A Dirichlet Multinomial Mixture model identified the existence of two distinct microbial rhinotypes, the first one characterized by significantly higher cotinine levels, lower alpha diversity, and enrichment of these taxa.

**Conclusion:** Children with higher cotinine levels had reduced alpha diversity and a distinct nasal rhinotype. Our results suggest TSE is associated with alterations of the nasal microbiome and identify a rhinotype as a potential biomarker for TSE.

## INTRODUCTION

The early life microbiome is critical for the establishment of immune responses, which can impact the risk of diseases later in life^1^. The nasal cavity can serve as a reservoir for microbes implicated in the pathogenesis of respiratory conditions^2-5^. Understanding factors such as secondhand tobacco smoke exposure (TSE) that shape microbial colonization in young children is therefore crucial to develop preventive approaches against respiratory conditions mediated by the nasal microbiome. Nicotine, a pro-inflammatory agent^6^ with selective anti-bacterial properties^7^, can influence the nasal and oral microbiome in adults^8,9^. Smokers have been found to carry more pathogenic strains in their nasal microbiomes than non-smokers^8-10^. The oral mucosa also has lower diversity in smokers compared to non-smokers^11^, and tobacco-associated COPD is associated with alterations in the nasal microbiome in adults^8^. Smoking can disrupt the adult nasal microbiome and adult-smoker caregivers can transfer pathogens to their children, including *Streptococcus pneumoniae*^12-14^, *Streptococcus pyogenes*^13^, *Staphylococcus aureus*^14^, and *Haemophilus influenzae* and *Moraxella catarrhalis*^13,14^. Furthermore, children exposed to tobacco smoke have lower gut microbiome diversity^15^.

Prior studies of conditions such as chronic rhinosinusitis, allergic rhinitis, asthma, and otitis media have also identified a disruption of the nasal and airway microbiome in both adults and children. For example, chronic rhinosinusitis is associated with higher levels of *Staphylococcus aureus* in infants^16^. In allergic rhinitis, the sinonasal microbiome of adults has higher diversity during the allergic season^16^, while children who reported tobacco smoke exposure had lower diversity in the anterior nostrils^17^. Early asthma diagnosis is associated with nasopharynx colonization of *Streptococcus* in infants^18^. In addition, pediatric and teenage nasal secretions dominated by *Streptococcus* species are not only associated with childhood asthma diagnosis, but with increased risk of rhinovirus infection^19^.

Importantly, these studies have not been performed in asymptomatic infants and children exposed to secondhand tobacco smoke. Further, caregiver reports of smoke exposure can underestimate actual exposure levels^20^. By contrast, cotinine, a metabolic breakdown of nicotine, is an established biomarker of TSE, and both salivary^21^ and urinary^22^ cotinine levels have been associated with pediatric respiratory hospitalization and disease. Here we investigate the impact of secondhand TSE, as quantified by cotinine in saliva and urine, on the pediatric nasal microbiome. We hypothesize that TSE levels will result in disruption of microbial communities, lower diversity and enrichment of specific taxa.

## METHODS

### Subject Data Collection

In this cross-sectional study, participants were recruited from New York and Maryland during December 2017 – May 2021. Recruitment included children from Mount Sinai Kravis Children’s Hospital (New York, NY; 6-month - 10-year-old children hospitalized for asthma or bronchiolitis, and children 1-2 years old or 4-5 years old presenting to the continuity clinic for a wellness visit), and the Johns Hopkins Children’s Center Bronchopulmonary Dysplasia Clinic (Baltimore, MD; 1-3-year-olds with a history bronchopulmonary dysplasia but stable at the time of enrollment presenting to the pulmonary outpatient clinic). Parents provided informed consent; children 7 and older provided assent. This study was approved by The Icahn School of Medicine at Mount Sinai Institutional Review Board (IRB STUDY-17-00125) and Johns Hopkins Institutional Review Board (IRB 00130419).

### Questionnaire data collection

A 43-question survey was administered to assess birth history (vaginal vs. caesarian birth), perinatal or other antibiotic use, other medication use, length of breastfeeding and family history of atopy. Tobacco exposure and demographic information were adapted from the Julius B. Richmond Center of Excellence Measurement Core standard survey questions^23^.

### TSE biomarkers

Salivary samples were collected using a cotton swab placed under the child’s tongue (when possible) or inside of the child’s mouth. The swabs were 125mm in length (SalivaBio’s Children’s Swab (SCS) System) to minimize any choking hazard. Once the swab was saturated (1-2 minutes), the sample of saliva was spun down and collected into a tube. Saliva in the tube was then frozen at -80C.

Urinary samples were collected for 1–2-year-old participants by placing cotton balls in the child’s diaper and waiting about one hour to obtain the sample. Cotton balls were placed in urine specimen cups. Within 30 minutes of collection, the cotton balls were placed in a syringe and the plunger was pressed to allow extraction of urine from the cotton balls into the specimen cup, which was then inverted 10 times. Urine was then pipetted into 2mL aliquots and immediately stored at -80C.

Saliva cotinine levels were measured using the Salimetrics Salivary Cotinine ELISA (#1-2112), with a limit of detection of 0.015 ng/mL^24^. Urinary cotinine was analyzed using a slightly modified method developed by Bernert et al^25^. After adding isotope labeled internal standards into 200 mL of saliva, samples were hydrolyzed with glucuronidase, purified using the supported liquid extraction method and finally concentrated in 100 mL water. A 5 mL aliquot of concentrated solution was injected into a UHPLC and analyzed using tandem mass spectrometry. The limit of detection for urinary cotinine was 0.030 ng/mL. A laboratory blank and quality control were simultaneously processed and analyzed for quality assessment.

### Microbiome characterization and analysis

The nasal microbiome was sampled using a standard DFA brush. The brush was placed in the participant nostril and swabbed around for up to 10 seconds per nostril for a total of 20 seconds. Samples were stored at -80C.

DNA was extracted using the Qiagen DNeasy PowerLyzer Powersoil Kit. Quantitation was performed on the Nanodrop 8000 Spectrophotometer. Library amplification was performed using Platinum Hot Start PCR 2x Master Mix from Invitrogen by Thermo Fisher Scientific using forward (515F) and reverse (806R) primers for the V4 region following the Earth Microbiome standard protocols^26,27,28^. DNA quantitation was performed using the Quant-iT Picogreen dsDNA Assay Kit from Invitrogen by Thermo Fisher Scientific. Sequencing cleanup was done using the QIAquick PCR Purification Kit followed by quantitation using the Qubit dsDNA HS Assay Kit on the Qubit 2.0 Fluorometer from Invitrogen by Life Technologies. Sequencing was performed at the NYU Genome Technology Center on an Illumina MiSeq Sequencing Instrument using 2x250bp read length.

Single barcode paired-end microbiome sequencing data was demultiplexed and processed using QIIME 2^29^. Reads were denoised using DADA2^30^ and mapped to GreenGenes for taxonomic assignment^31^. Rarefaction was performed at 3,000 reads per sample. Three experimental controls from elution buffers were included in the sequencing run. Two blank swabs were tested in an additional sequencing run to rule out the possibility of contamination from the swabs directly. Alpha diversity was measured using Shannon entropy, and beta diversity was estimated using Bray-Curtis distances. Differential abundance analysis was performed using LEfSe 1.1.01^32^. Taxa that could not be identified below the order level were removed, with the exception of those that were the only representative of a given phylum and not a known contaminant^33^. Known contaminants were removed from the final taxa table^33^. The list of taxa identified and filtered are reported in **Supplementary Table 1 and Table 2** respectively.

For those taxa that could not be fully identified at the species level, we performed BLAST searches against NCBI, selecting the top hit as suggestive of species-level identification. Because the bacteria Dolosigranulum can be misclassified as Alloiococcus^34^, we confirmed through a BLAST search that Alloicoccus sequences in our dataset in fact represent Dolosigranulum.

### Microbiome Co-occurrence Network

Network co-occurrence analysis was performed using SparCC with default parameters and using genus-level taxa after removing contaminants^35^. Nodes of the network represent bacterial genera and edges represent significant correlations between nodes, with shorter edges indicating stronger correlations. Networks were visualized with Cytoscape v3.9.1^36^, using the edge weighted spring embedded layout and with edge lengths representing the strength of the correlation between taxa as estimated by SparCC.

### Dirichlet Multinomial Mixture Model

We used a Dirichlet Multinomial Mixture (DMM) model to infer the optimal number of microbiome community types (minimal number of clusters that maximize the amount of information in the data) that best describe our cohort. We used three different criteria to identify the number of clusters: Laplace, Akaike’s Information Criterion (AIC) and the Bayesian Information Criterion (BIC). The DMM model was generated using the packages *microbiome* 1.19.1 and *DirichletMultinomial* 1.36.0 in R 4.1.0^37^.

### Statistical analyses

Additional statistical analyses were conducted using R version 4.2.2. Comparison of continuous variables was done using the Mann-Whitney U-test. Correlation between salivary and urinary cotinine was done using the squared Pearson correlation coefficient. All tests were two-sided and P values less than 0.05 were considered statistically significant.

## RESULTS

### Cohort description

Out of 236 patients recruited, microbiome samples were obtained from 221 of them. Four samples had low sequencing coverage and were discarded, resulting in a total of 217 microbiome samples from patients. After cotinine and microbiome sample processing, our cohort included data from 181 inpatient and outpatient children recruited at Mount Sinai Kravis Children’s Hospital and 36 outpatient children from Johns Hopkins Children’s Center. **Table 1** describes the demographics of the enrolled subjects. Study participants were classified into four categories: inpatients, outpatients with a history of bronchopulmonary dysplasia (hereafter “BPD”; note that these patients were otherwise healthy at time of enrollment), healthy outpatients between the ages of 1-2 years old, and healthy outpatients between the ages of 4-5 years old. Salivary cotinine data was obtained for 183 participants, and urinary cotinine for 64 participants, with 44 participants having both salivary and urinary cotinine data (**Figure 1**).

**Table 1.**
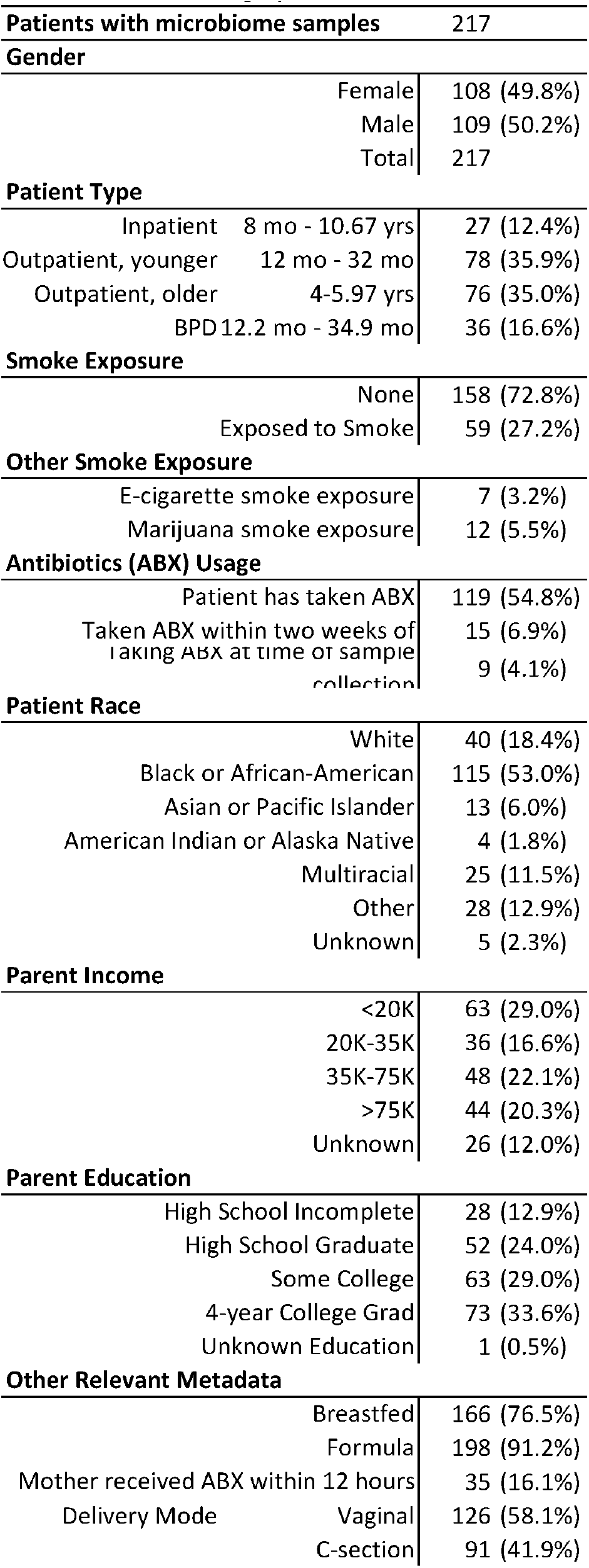
Cohort demographics.

**Figure 1.**
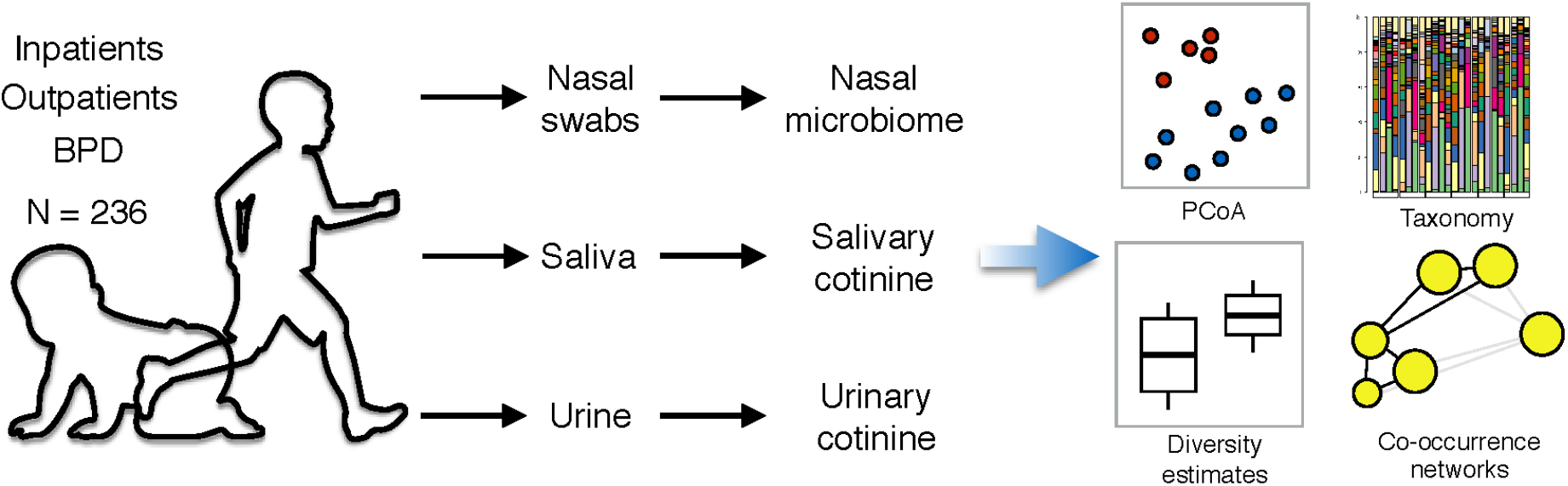
Study design, assays and analytical approaches.

### Nasal microbiome across age, patient type and collection site

The taxonomic profiles of children in our cohort were dominated by three genera, Moraxella, Corynebacterium and Dolosigranulum, although with high interpersonal variability (**Figure 2A**, b). Alpha diversity was not significantly different between collection sites (U-test, p = 0.058) (**Supplementary Figure 1A, Table 3**) or patient types (**Supplementary Figure 1B**). Principal Coordinate Analysis (PCoA) based on Bray-Curtis distances did not reveal a clear clustering structure associated with collection site (**Figure 2B**) although PERMANOVA analysis found a significant difference by site (p = 0.001). Participants did not cluster visually in PCoA based on patient type (**Figure 2C**), although there was a statistically significant difference between BPD outpatients and all other patient groups (**Supplementary Table 3**).

**Figure 2.**
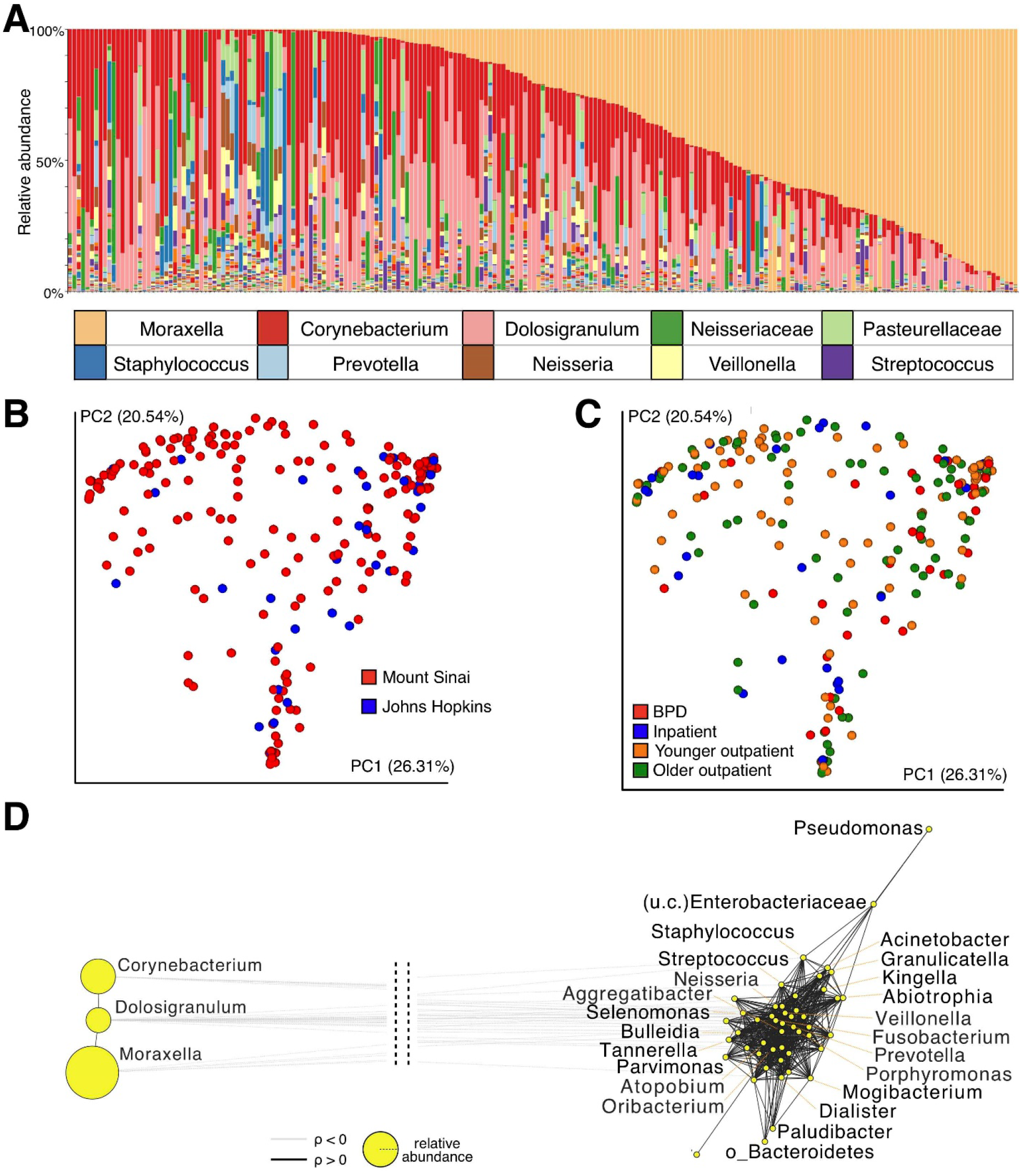
Microbiome sample description. **A**. Relative abundance across all samples (n=217), sorted by most abundant taxa. Top 10 most abundant taxa described in the figure, full list available in Supplementary Table 1. **B**. Principal coordinates plot based on Bray-Curtis distances colored by site of sample collection. Johns Hopkins Hospital: blue; Mount Sinai Hospital: red. **C**. Principal coordinates plot based on Bray-Curtis distances colored by subject. Bronchopulmonary Dysplasia (BPD) outpatients: red; asthma inpatients: blue; bronchiolitis inpatients: orange, younger (1-2-year-old) outpatients: purple; older (4-5-year-old) outpatients: green. **D**. Co-occurrence network estimated using SparCC with default parameters. Nodes represent bacterial taxa, with size of the node representing relative abundance. Edges indicate significant correlations (black: positive correlation; gray: negative) between taxa, with length of the edge being inversely proportional to correlation strength (shorter edges: stronger correlation).

We investigated whether factors generally associated with early-life microbiome composition, such as antibiotic use, feeding mode or delivery mode, were associated with microbial diversity estimates. None were found to correlate significantly with cotinine levels or microbiome diversity, including patient age (**Supplementary Figure 2**), except for feeding mode, where infants that only received formula had more diverse microbiomes than those who received only breastmilk or a combination of the two (Mann-Whitney, MW p = 0.026) (**Supplementary Figure 3**). Overall, these results suggest that several factors that impact the microbiome in other body sites do not have a significant effect on nasal bacterial communities in our cohort.

### Network analysis identifies cliques of co-occurring taxa

We found a large separation between two clusters of distinct taxa (**Figure 2D**): the three most abundant taxa in the cohort (Corynebacterium, Dolosigranulum, Moraxella) were strongly and positively correlated between themselves. Low abundance taxa, on the other hand, clustered separately, with weak positive correlations between themselves and strong negative correlations with the more abundant taxa. Overall, these results suggest that the nasal microbiome of children in our cohort is configured either as dominated by a small number of high abundance taxa or by a large number of low abundance taxa.

### Cotinine levels are associated with a distinct nasal microbial profile

Salivary and urinary cotinine levels were significantly correlated (**Supplementary Figure 4**, p=1.123e-7). Higher salivary cotinine (**Figure 3A**) was generally associated with lower alpha diversity, and higher urinary cotinine showed a similar trend (**Figure 3B**). Patients with salivary cotinine less than 1.5 ng/µL, the threshold for TSE exposure in saliva^38^, had significantly higher diversity compared to patients with levels equal or greater than 9.5ng/ µL (p=0.042, Mann-Whitney U-test). The effect was not statistically significant when comparing patients with urinary cotinine less than 2 ng/µL, the threshold for high TSE in urine^39^, to patients with levels equal or greater than 10 ng/µL.

**Figure 3.**
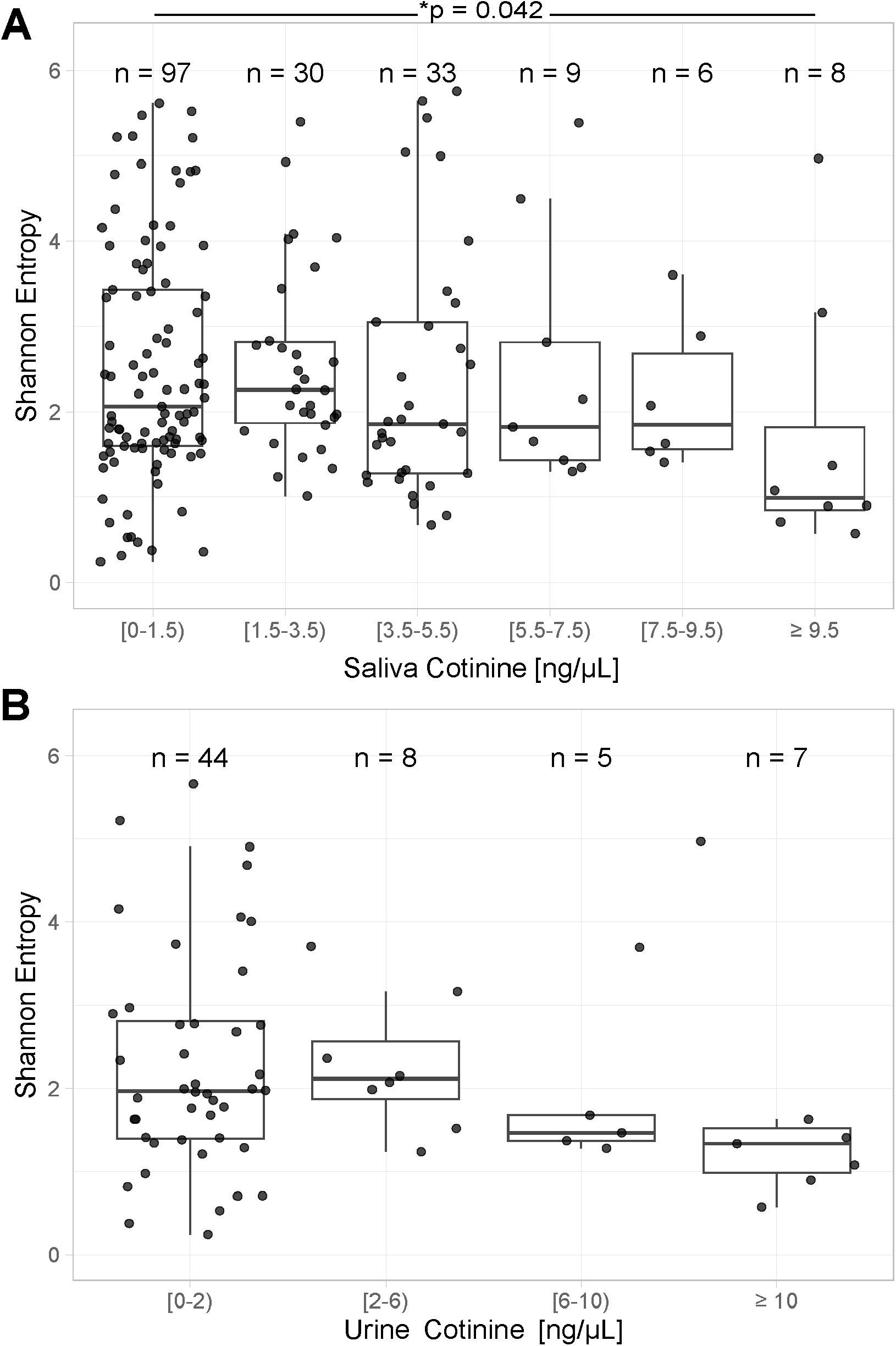
Diversity analysis identifies bacteria associated with cotinine levels in saliva. **A**. Distribution of alpha diversity values (Shannon entropy) at increasing levels of salivary cotinine (n=183). Cotinine data is binned in intervals of 2 ng/µL, except for the first bin, which includes patients with levels less than 1.5 ng/µL. **B**. Distribution of alpha diversity values (Shannon entropy) at increasing levels of urinary cotinine (n=64). Cotinine data is binned in intervals of 4 ng/µL, except for the first bin which includes patients with levels less than 2 ng/µL.

### Dirichlet Multinomial Mixture reveals two distinct microbial rhinotypes associated with cotinine

Using DMM, we identified two microbiome community types (“rhinotypes”) in our cohort (**Figure 4A**). Rhinotype 1 was primarily characterized by three genera – Moraxella, Dolosigranulum and Corynebacterium, which had higher relative abundance than in rhinotype 2 (**Supplementary Figure 5**). Rhinotype 2 was dominated by a mixture of different bacteria, including numerous taxa with low relative abundance (**Figure 4B**). Nasal rhinotypes significantly separated subjects based on Bray-Curtis distances (**Figure 4C**; p=0.001 PERMANOVA). Notably, rhinotype 1 children had significantly lower alpha diversity than rhinotype 2 children (MW, p<2.2 e-16) (**Figure 4D**). This effect was not related to whether the patient was currently on antibiotics (MW p = 0.64), had taken antibiotics in the last week (MW p = 0.94), or had ever taken antibiotics (MW, p=0.62). Furthermore, rhinotype 1 was also associated with significantly higher salivary cotinine levels (MW, p=0.0219) (**Figure 4E**). Overall, these results demonstrate that children in our cohort can be classified into two distinct nasal rhinotypes which are associated with significant differences in microbial composition, diversity and salivary cotinine levels.

**Figure 4.**
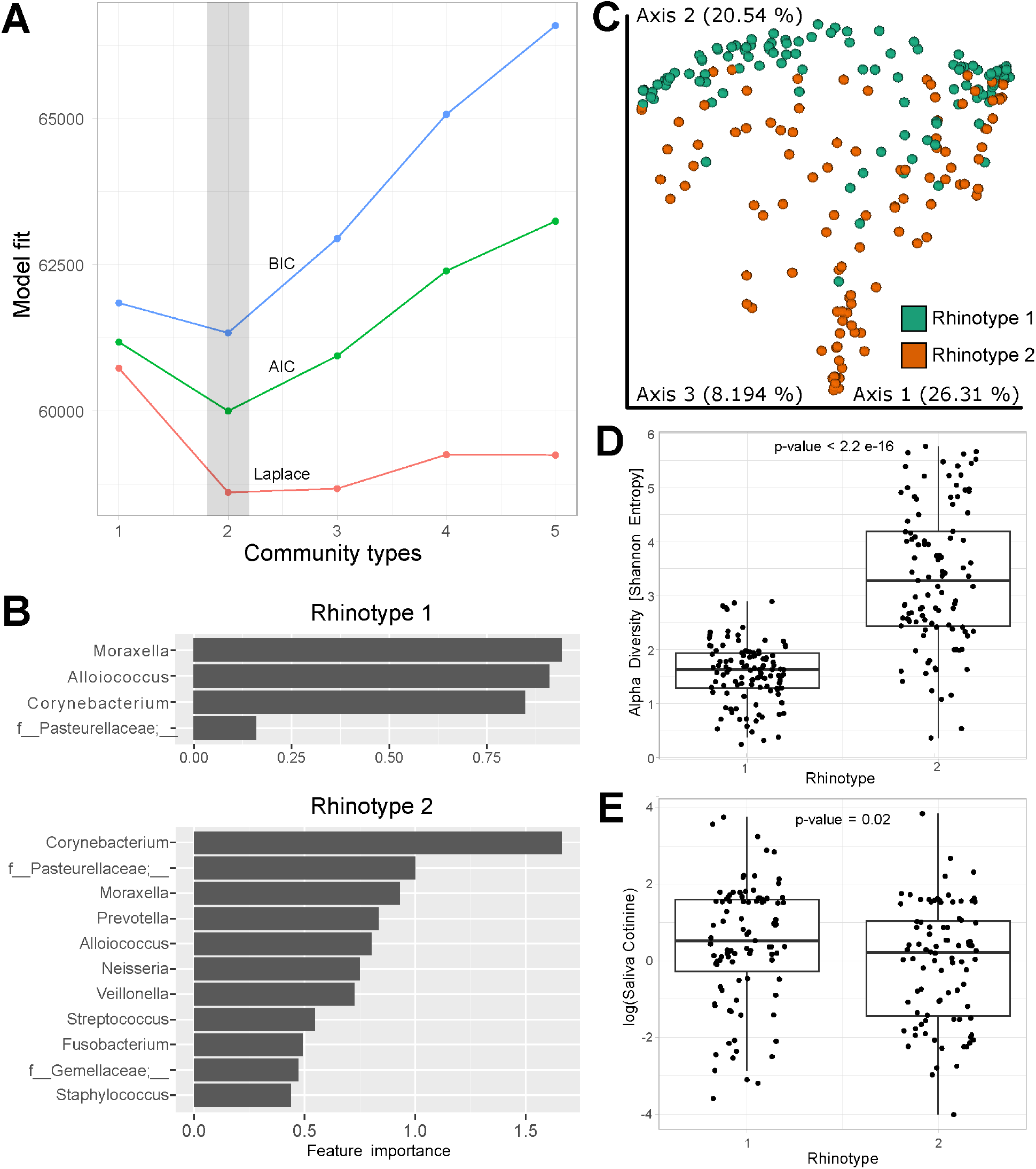
Dirichlet Multinomial Mixture (DMM) model reveals two rhinotypes with significant differences in microbiome and salivary cotinine profiles. **A**. Model fit with three different criteria (Laplace, Akaike information criterion, Bayesian information criterion) identifies two as the optimal number of nasal microbiome clusters (rhinotypes 1 and 2) (n=217). **B**. Features characteristic of each rhinotype predicted by the DMM model. **C**. Principal coordinate analysis plot, with samples colored by rhinotype. **D**. Alpha diversity in each rhinotype (n=217, p<0.001). **E**. Saliva cotinine levels in each rhinotype (n=183, p=0.02). Significance calculated with the Mann-Whitney U-test.

## DISCUSSION

In this work, we examined the nasal microbiome of children and its relation to salivary and urinary cotinine levels reflecting tobacco smoke exposure. We found an association between higher cotinine levels and a distinct microbiome rhinotype with low diversity. In addition, we observed that inpatients had higher urinary cotinine compared to younger outpatients (p=0.005 by U-test), suggesting that changes in the nasal microbiome induced by TSE could contribute to the higher risk of respiratory morbidities in children exposed to secondhand smoke. To our knowledge, our study is the largest investigation to date of the anterior nares microbiome in a pediatric cohort with cotinine TSE data. McCauley et al used nasal secretion samples from 413 subjects aged 6-17^19^, while most of our subjects are younger than 6 years. Using data from the Childhood Asthma Study, Teo and colleagues studied 234 infants less than 1 year old, but using nasopharynx samples^18^. Neither of those studies included data on TSE as measured by salivary or urinary cotinine.

In contrast with the maturation of the gut microbiome^40^, we did not observe changes with age in the nasal microbiome diversity in our cohort (**Supplementary Figure 2**). On the other hand, infants exclusively fed formula had higher microbiome diversity (**Supplementary Figure 3**), suggesting that feeding mode may influence the development of the nasal microbiome, similar to previous studies in the upper respiratory tract microbiome^41^ and gut microbiome^41,42^. The most prevalent bacteria found in our cohort (**Figure 2A**) are in concordance with prior characterizations of the nasal microbiome^16,43^, as well as with culture-based studies of the pediatric nasopharynx microbiome^12,14,44^. Propionibacerium, Flavobacterium, Haemophilus and Dolosigranulum have also been reported as part of the microbiome in nasal secretions^19^, the anterior nares^16^ and the nasopharynx^18,34^.

We found that high levels of salivary or urinary cotinine were associated with lower nasal microbiome diversity in our cohort (**Figure 3**). A prior study demonstrated that infants exposed to secondhand tobacco smoke, as measured by urinary cotinine levels, had lower gut microbiome diversity^15^. As nicotine is a toxin, it is plausible that it exerts selective anti-microbial effects^7^. Importantly, our unbiased approach to identify two distinct nasal rhinotypes confirms this observation, as the rhinotype with higher cotinine levels had significantly lower diversity and distinct composition (**Figure 4**). Previous studies have also categorized subjects based on their microbiome to identify clinical phenotypes in the lung^45^, gut^46^ or the vagina^47^. To our knowledge, this is the first time that this approach is used to characterize the nasal microbiome of a pediatric cohort, and our results suggest that rhinotypes could be used to stratify subjects based on their levels of exposure to tobacco smokel.

There are limitations to our study. The data was cross-sectional and cannot establish whether the observed changes in the nasal microbiome are the result of recent acute smoke exposure or a gradual adaptation due to prolonged exposure. In addition, there is the possibility of recall bias in our assessments of other potential factors that could alter the microbiome.

Overall, our findings demonstrate that children exposed to high cotinine levels have a distinct microbial rhinotype with lower microbial diversity and high abundance of Moraxella, Dolosigranulum and Corynebacterium. These results suggest a disruption in the nasal microbiome community among children exposed to tobacco smoke. While continued efforts to support tobacco cessation among parents of young children are critical, there is also a strong need to further investigate how the microbiome of children exposed to tobacco smoke evolves over time, and how such changes impact host immune responses and the risk of respiratory conditions.

## DATA AVAILABILITY STATEMENT

Sequencing data is publicly available through NCBI as BioProject PRJNA993245.

## FIGURES and TABLES

**Supplementary Figure 1. Shannon Entropy by collection site and patient type. A**. Shannon entropy alpha diversity separated by site. Mann-Whitney U-test p-value 0.058, (n=217). **B**. Shannon entropy alpha diversity separated by patient type. Mann-Whitney U-test p-values are described in Supplementary Table 3 (n=217).

**Supplementary Figure 2. Alpha and Beta Diversity as a function of age. A**. Shannon entropy as a function of age (n=217). Line represents loess smoothing applied to the dataset. Gray region represents 95% confidence interval. **B**. Shannon entropy as a function of age (n=217). Line represents linear model fit applied to the dataset. Grey region represents 95% confidence interval. **C**. PCoA analysis based on Bray-Curtis first principal component by patient age. Line represents LOESS interpolation, with 95% confidence interval indicated by grey shading. **D**. PCoA analysis based on Bray-Curtis first principal component by patient age. Line represents linear model fit applied to the dataset with 95% confidence interval indicated by grey shading.

**Supplementary Figure 3. Nasal alpha diversity is higher in formula-fed infants**. ‘Yes’ indicates patients were only given formula. ‘No’ indicates patients given either breastmilk only or a combination of breastmilk and formula. Data colored by subject age in months. Shannon entropy p-value 0.026, by Mann-Whitney U-test.

**Supplementary Figure 4. Correlation between salivary and urinary cotinine levels**. Pearson correlation for salivary vs urinary cotinine levels for all patients in which both were collected at the time of sampling.

**Supplementary Figure 5. Enrichment of specific taxa in each rhinotype. A**. Dolosigranulum abundance is higher in rhinotype 1 infants (MW U-test p-value = 4.5e-6). **B**. Corynebacterium abundance is higher in rhinotype 1 infants, although not significantly (MW U-test p-value = 0.055). **C**. Moraxella abundance is higher in rhinotype 1 infants (MW U-test p-value = 7.9e-7).

**Supplementary Table 1**. List of all taxa in the dataset (non-contaminants)

**Supplementary Table 2**. All contaminant taxa that were removed

**Supplementary Table 3**. Comparisons between patient types for alpha (Mann-Whitney U-test) and beta diversity (PERMANOVA). Significant differences indicated in boldface.

## REFERENCES

1 Tamburini, S., Shen, N., Wu, H. C. & Clemente, J. C. The microbiome in early life: implications for health outcomes. Nat Med 22, 713–722, doi:10.1038/nm.4142 (2016).

2 Jochems, S. P., Ferreira, D. M. & Smits, H. H. Microbiota and compartment matter in the COVID-19 response. Nat Immunol 22, 1350–1352, doi:10.1038/s41590-021-01041-w (2021).

3 Toivonen, L. et al. Early nasal microbiota and acute respiratory infections during the first years of life. Thorax 74, 592–599, doi:10.1136/thoraxjnl-2018-212629 (2019).

4 van den Munckhof, E. H. A. et al. Nasal microbiota dominated by Moraxella spp. is associated with respiratory health in the elderly population: a case control study. Respir Res 21, 181, doi:10.1186/s12931-020-01443-8 (2020).

5 Aurora, R. et al. Contrasting the microbiomes from healthy volunteers and patients with chronic rhinosinusitis. JAMA Otolaryngol Head Neck Surg 139, 1328–1338, doi:10.1001/jamaoto.2013.5465 (2013).

6 Zhang, W. et al. Nicotine in Inflammatory Diseases: Anti-Inflammatory and Pro-Inflammatory Effects. Front Immunol 13, 826889, doi:10.3389/fimmu.2022.826889 (2022).

7 Cussotto, S., Clarke, G., Dinan, T. G. & Cryan, J. F. Psychotropics and the Microbiome: a Chamber of Secrets. Psychopharmacology (Berl) 236, 1411–1432, doi:10.1007/s00213-019-5185-8 (2019).

8 Agarwal, D. M. et al. Disruptions in oral and nasal microbiota in biomass and tobacco smoke associated chronic obstructive pulmonary disease. Arch Microbiol 203, 2087–2099, doi:10.1007/s00203-020-02155-9 (2021).

9 Pfeiffer, S. et al. Different responses of the oral, nasal and lung microbiomes to cigarette smoke. Thorax 77, 191–195, doi:10.1136/thoraxjnl-2020-216153 (2022).

10 Charlson, E. S. et al. Disordered microbial communities in the upper respiratory tract of cigarette smokers. PLoS One 5, e15216, doi:10.1371/journal.pone.0015216 (2010).

11 Yu, G. et al. The effect of cigarette smoking on the oral and nasal microbiota. Microbiome 5, 3, doi:10.1186/s40168-016-0226-6 (2017).

12 Greenberg, D. et al. The contribution of smoking and exposure to tobacco smoke to Streptococcus pneumoniae and Haemophilus influenzae carriage in children and their mothers. Clin Infect Dis 42, 897–903, doi:10.1086/500935 (2006).

13 Brook, I. Effects of exposure to smoking on the microbial flora of children and their parents. Int J Pediatr Otorhinolaryngol 74, 447–450, doi:10.1016/j.ijporl.2010.01.006 (2010).

14 Bugova, G., Janickova, M., Uhliarova, B., Babela, R. & Jesenak, M. The effect of passive smoking on bacterial colonisation of the upper airways and selected laboratory parameters in children. Acta Otorhinolaryngol Ital 38, 431–438, doi:10.14639/0392-100X-1573 (2018).

15 Northrup, T. F. et al. Thirdhand smoke associations with the gut microbiomes of infants admitted to a neonatal intensive care unit: An observational study. Environ Res 197, 111180, doi:10.1016/j.envres.2021.111180 (2021).

16 Rawls, M. & Ellis, A. K. The microbiome of the nose. Ann Allergy Asthma Immunol 122, 17–24, doi:10.1016/j.anai.2018.05.009 (2019).

17 Brindisi, G. et al. Allergic rhinitis, microbiota and passive smoke in children: A pilot study. Pediatr Allergy Immunol 33 Suppl 27, 22–26, doi:10.1111/pai.13621 (2022).

18 Teo, S. M. et al. The infant nasopharyngeal microbiome impacts severity of lower respiratory infection and risk of asthma development. Cell Host Microbe 17, 704–715, doi:10.1016/j.chom.2015.03.008 (2015).

19 McCauley, K. et al. Distinct nasal airway bacterial microbiotas differentially relate to exacerbation in pediatric patients with asthma. J Allergy Clin Immunol 144, 1187–1197, doi:10.1016/j.jaci.2019.05.035 (2019).

20 Collaco, J. M. et al. Hair nicotine levels in children with bronchopulmonary dysplasia. Pediatrics 135, e678–686, doi:10.1542/peds.2014-2501 (2015).

21 Howrylak, J. A. et al. Cotinine in children admitted for asthma and readmission. Pediatrics 133, e355–362, doi:10.1542/peds.2013-2422 (2014).

22 Tovar, M. F. et al. Prevalence of urinary cotinine levels in children under 5 years of age during consultations for acute respiratory disease at the emergency department of the Universidad de La Sabana clinic. BMC Pediatr 20, 296, doi:10.1186/s12887-020-02193-8 (2020).

23 Pediatrics, A. A. o. Measurement Core, <https://www.aap.org/en/patient-care/tobacco-control-and-prevention/tobacco-control-research/measurement-core/> (

24 Granger, D. A. et al. Individual differences in salivary cortisol and alpha-amylase in mothers and their infants: relation to tobacco smoke exposure. Dev Psychobiol 49, 692–701, doi:10.1002/dev.20247 (2007).

25 Bernert, J. T. et al. Urinary tobacco-specific nitrosamines and 4-aminobiphenyl hemoglobin adducts measured in smokers of either regular or light cigarettes. Nicotine Tob Res 7, 729–738, doi:10.1080/14622200500259762 (2005).

26 Caporaso, J. G. et al. Ultra-high-throughput microbial community analysis on the Illumina HiSeq and MiSeq platforms. ISME J 6, 1621–1624, doi:10.1038/ismej.2012.8 (2012).

27 Caporaso, J. G. et al. Global patterns of 16S rRNA diversity at a depth of millions of sequences per sample. Proc Natl Acad Sci U S A 108 Suppl 1, 4516–4522, doi:10.1073/pnas.1000080107 (2011).

28 Thompson, L. R. et al. A communal catalogue reveals Earth’s multiscale microbial diversity. Nature 551, 457–463, doi:10.1038/nature24621 (2017).

29 Bolyen, E. et al. Reproducible, interactive, scalable and extensible microbiome data science using QIIME 2. Nat Biotechnol 37, 852–857, doi:10.1038/s41587-019-0209-9 (2019).

30 Callahan, B. J. et al. DADA2: High-resolution sample inference from Illumina amplicon data. Nat Methods 13, 581–583, doi:10.1038/nmeth.3869 (2016).

31 McDonald, D. et al. An improved Greengenes taxonomy with explicit ranks for ecological and evolutionary analyses of bacteria and archaea. Isme J 6, 610–618, doi:10.1038/ismej.2011.139 (2012).

32 Segata, N. et al. Metagenomic biomarker discovery and explanation. Genome Biol 12, R60, doi:10.1186/gb-2011-12-6-r60 (2011).

33 Salter, S. J. et al. Reagent and laboratory contamination can critically impact sequence-based microbiome analyses. BMC Biol 12, 87, doi:10.1186/s12915-014-0087-z (2014).

34 Lappan, R. et al. A microbiome case-control study of recurrent acute otitis media identified potentially protective bacterial genera. BMC Microbiol 18, 13, doi:10.1186/s12866-018-1154-3 (2018).

35 Friedman, J. & Alm, E. J. Inferring correlation networks from genomic survey data. PLoS Comput Biol 8, e1002687, doi:10.1371/journal.pcbi.1002687 (2012).

36 Shannon, P. et al. Cytoscape: a software environment for integrated models of biomolecular interaction networks. Genome Res 13, 2498–2504, doi:10.1101/gr.1239303 (2003).

37 Holmes, I., Harris, K. & Quince, C. Dirichlet multinomial mixtures: generative models for microbial metagenomics. PLoS One 7, e30126, doi:10.1371/journal.pone.0030126 (2012).

38 Polanska, K. et al. Estimation of Saliva Cotinine Cut-Off Points for Active and Passive Smoking during Pregnancy-Polish Mother and Child Cohort (REPRO_PL). Int J Environ Res Public Health 13, doi:10.3390/ijerph13121216 (2016).

39 Sangmo, L. et al. Secondhand marijuana exposure in a convenience sample of young children in New York City. Pediatr Res 89, 905–910, doi:10.1038/s41390-020-0958-7 (2021).

40 Yatsunenko, T. et al. Human gut microbiome viewed across age and geography. Nature 486, 222–227, doi:10.1038/nature11053 (2012).

41 Rosas-Salazar, C. et al. Exclusive breast-feeding, the early-life microbiome and immune response, and common childhood respiratory illnesses. J Allergy Clin Immunol 150, 612–621, doi:10.1016/j.jaci.2022.02.023 (2022).

42 Ma, J. et al. Comparison of the Gut Microbiota in Healthy Infants With Different Delivery Modes and Feeding Types: A Cohort Study. Front Microbiol 13, 868227, doi:10.3389/fmicb.2022.868227 (2022).

43 Bender, M. E. et al. A Comparison of the Bacterial Nasal Microbiome in Allergic Rhinitis Patients Before and After Immunotherapy. Laryngoscope 130, E882–E888, doi:10.1002/lary.28599 (2020).

44 Brook, I. & Gober, A. E. Recovery of potential pathogens in the nasopharynx of healthy and otitis media-prone children and their smoking and nonsmoking parents. Ann Otol Rhinol Laryngol 117, 727–730, doi:10.1177/000348940811701003 (2008).

45 Segal, L. N. et al. Enrichment of the lung microbiome with oral taxa is associated with lung inflammation of a Th17 phenotype. Nat Microbiol 1, 16031, doi:10.1038/nmicrobiol.2016.31 (2016).

46 Costea, P. I. et al. Enterotypes in the landscape of gut microbial community composition. Nat Microbiol 3, 8–16, doi:10.1038/s41564-017-0072-8 (2018).

47 France, M. T. et al. VALENCIA: a nearest centroid classification method for vaginal microbial communities based on composition. Microbiome 8, 166, doi:10.1186/s40168-020-00934-6 (2020).

