## Supplemental Information for "Tobacco Smoke Exposure is Characterized by a Distinct Nasal Rhinotype in a Pediatric Population"

#### Supplemental Material

### Tobacco Smoke Exposure is Associated with a Distinct Nasal Rhinotype and Respiratory Illnesses in a Pediatric Population

**Supplementary Figure 1:** Shannon Entropy by site and patient type.

**Supplementary Figure 2:** Alpha and Beta Diversity as a function of age.

**Supplementary Figure 3:** Nasal microbiome alpha diversity is higher in formula-fed infants.

**Supplementary Figure 4:** The correlation between salivary cotinine and urinary cotinine levels.

**Supplementary Figure 5:** Enrichment of specific taxa in each rhinotype.

**This supplemental material has been provided by the authors to give readers additional information about their work.**

**Supplementary Figure 1:** Shannon Entropy by site and patient type.

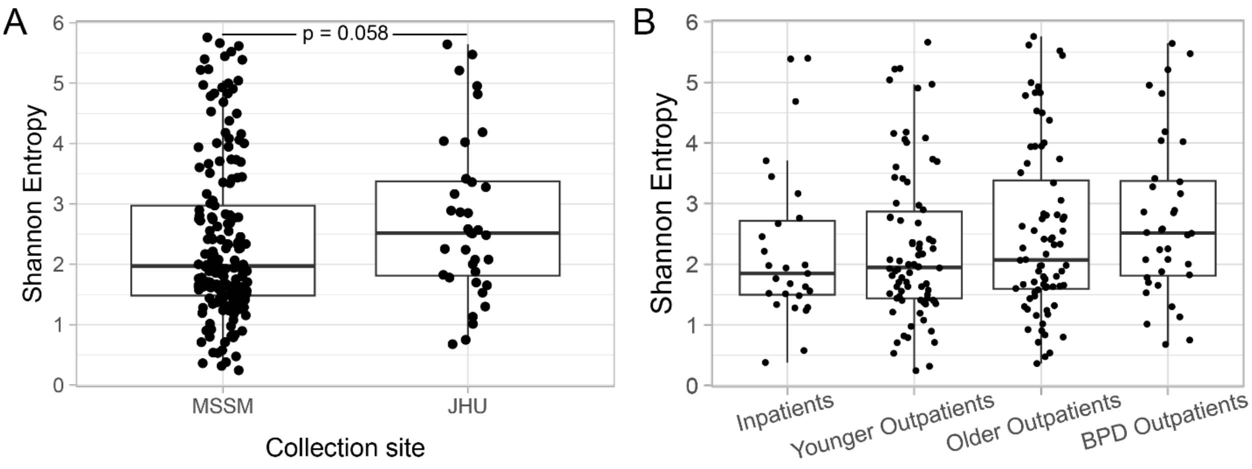

**Supplementary Figure 2:** Alpha and Beta Diversity as a function of age.

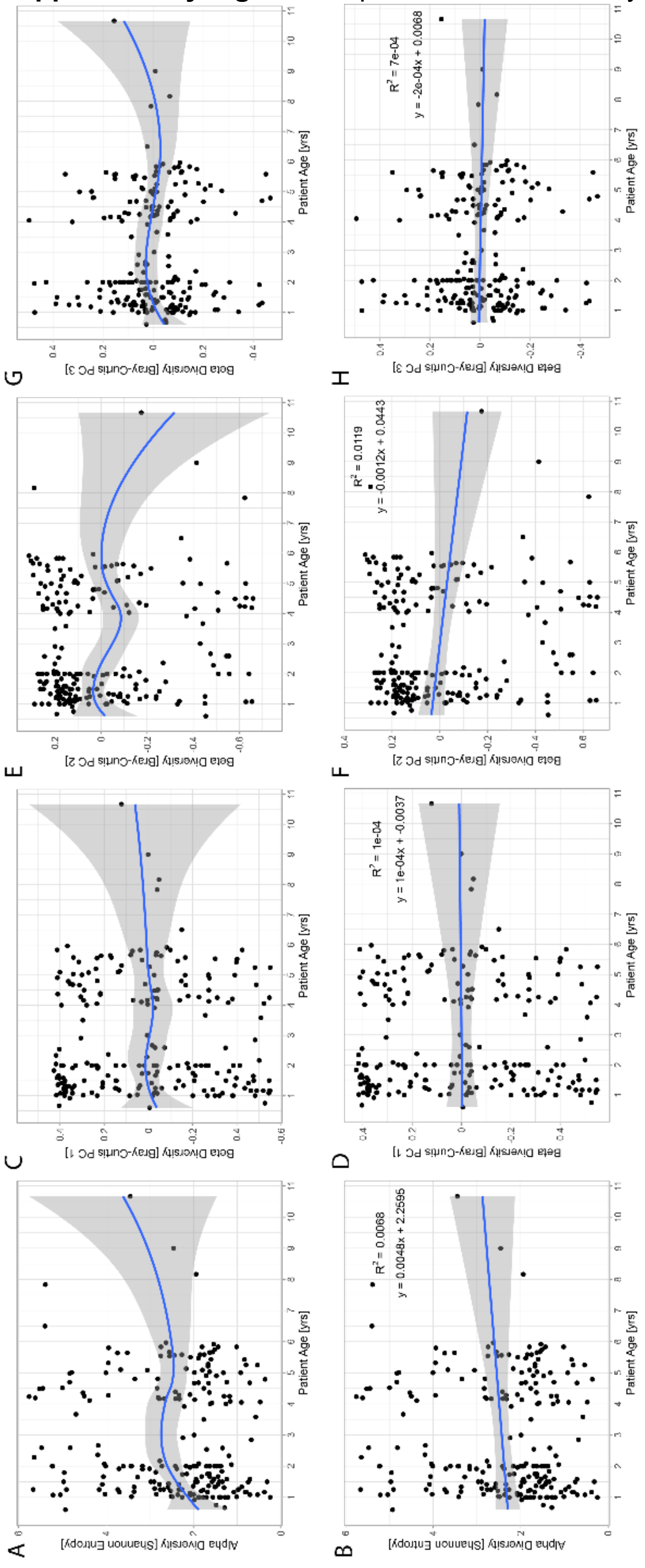

**Supplementary Figure 3:** Nasal microbiome alpha diversity is higher in formula-fed infants.

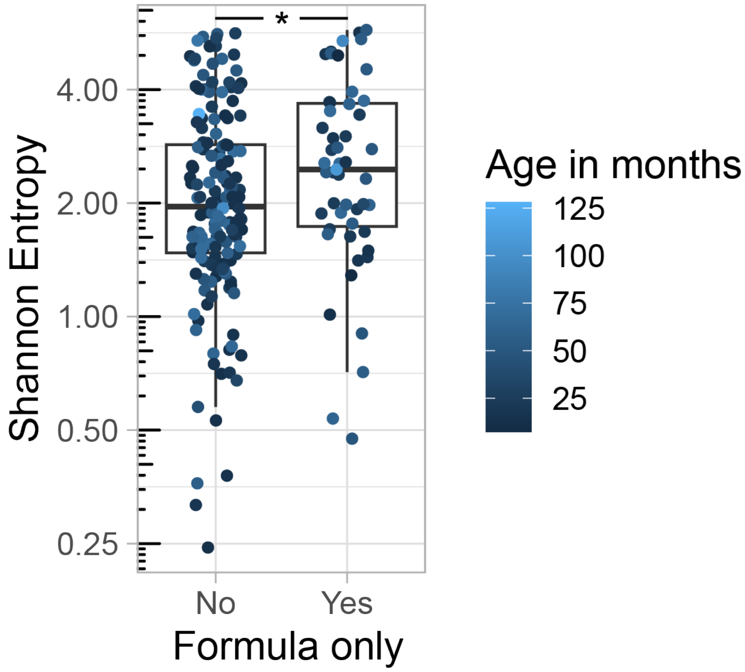


**Supplementary Figure 4:** The correlation between salivary cotinine and urinary cotinine levels.

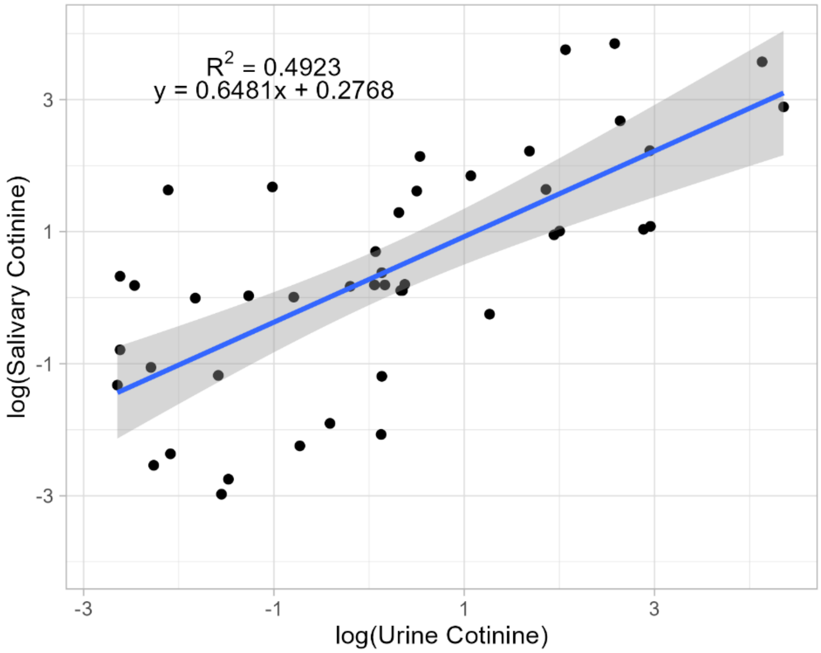

**Supplementary Figure 5:** Enrichment of specific taxa in each rhinotype.

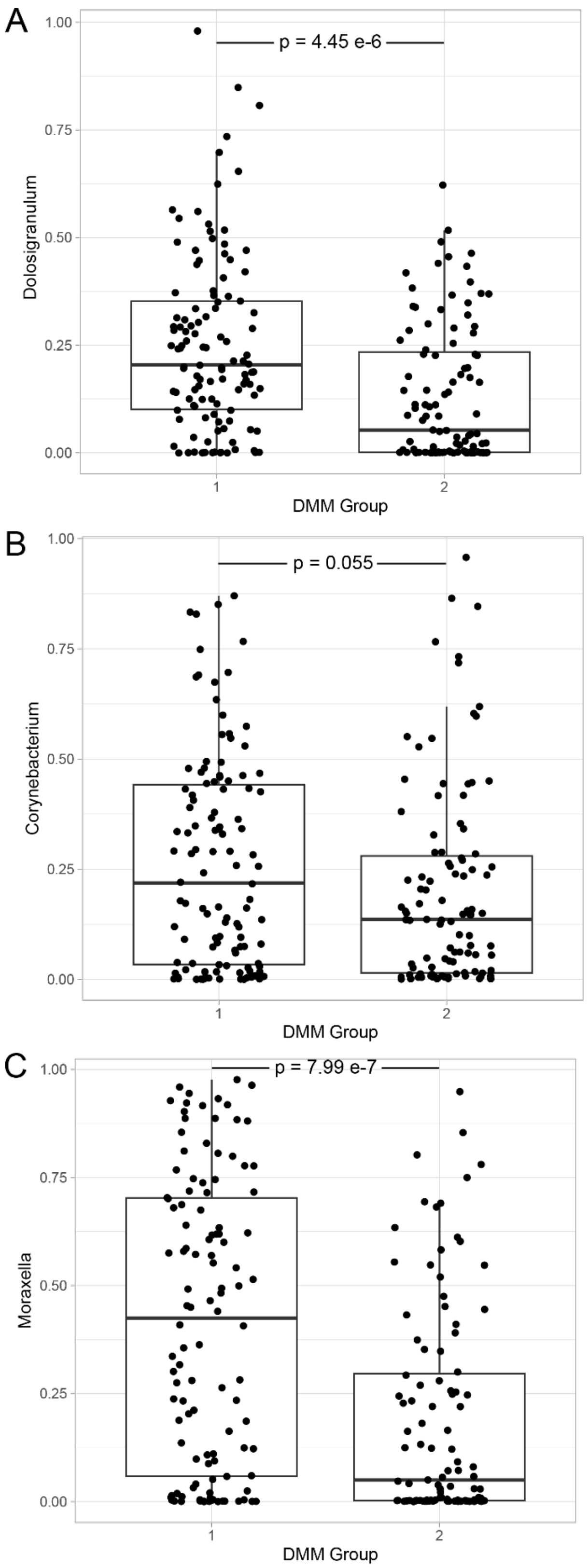
