## Supplemental Tables 1-3 for "Tobacco Smoke Exposure is Characterized by a Distinct Nasal Rhinotype in a Pediatric Population"

**eTable 1**

Unassigned;__;__;__;__;__
k__Archaea;p__Euryarchaeota;c__Methanobacteria;o__Methanobacteriales;f__Methanobacteriaceae;g__Methanobrevibacter k__Archaea;p__Euryarchaeota;c__Methanobacteria;o__Methanobacteriales;f__Methanobacteriaceae;g__Methanosphaera k__Bacteria;__;__;__;__;__
k__Bacteria;p__Acidobacteria;c__Acidobacteria-6;o__iii1-15;f__;g__
k__Bacteria;p__Acidobacteria;c__[Chloracidobacteria];o__RB41;f__Ellin6075;g__ k__Bacteria;p__Actinobacteria;c__Actinobacteria;o__Actinomycetales;__;__
k__Bacteria;p__Actinobacteria;c__Actinobacteria;o__Actinomycetales;f__Actinomycetaceae;g__ k__Bacteria;p__Actinobacteria;c__Actinobacteria;o__Actinomycetales;f__Actinomycetaceae;g__Actinobaculum k__Bacteria;p__Actinobacteria;c__Actinobacteria;o__Actinomycetales;f__Actinomycetaceae;g__Actinomyces k__Bacteria;p__Actinobacteria;c__Actinobacteria;o__Actinomycetales;f__Actinomycetaceae;g__Arcanobacterium k__Bacteria;p__Actinobacteria;c__Actinobacteria;o__Actinomycetales;f__Actinomycetaceae;g__Mobiluncus k__Bacteria;p__Actinobacteria;c__Actinobacteria;o__Actinomycetales;f__Actinomycetaceae;g__N09 k__Bacteria;p__Actinobacteria;c__Actinobacteria;o__Actinomycetales;f__Actinomycetaceae;g__Varibaculum k__Bacteria;p__Actinobacteria;c__Actinobacteria;o__Actinomycetales;f__Brevibacteriaceae;g__Brevibacterium k__Bacteria;p__Actinobacteria;c__Actinobacteria;o__Actinomycetales;f__Corynebacteriaceae;g__Corynebacterium k__Bacteria;p__Actinobacteria;c__Actinobacteria;o__Actinomycetales;f__Dermabacteraceae;g__ k__Bacteria;p__Actinobacteria;c__Actinobacteria;o__Actinomycetales;f__Dermabacteraceae;g__Brachybacterium k__Bacteria;p__Actinobacteria;c__Actinobacteria;o__Actinomycetales;f__Dermabacteraceae;g__Dermabacter k__Bacteria;p__Actinobacteria;c__Actinobacteria;o__Actinomycetales;f__Dermacoccaceae;g__Dermacoccus k__Bacteria;p__Actinobacteria;c__Actinobacteria;o__Actinomycetales;f__Dermatophilaceae;__ k__Bacteria;p__Actinobacteria;c__Actinobacteria;o__Actinomycetales;f__Dietziaceae;__ k__Bacteria;p__Actinobacteria;c__Actinobacteria;o__Actinomycetales;f__Dietziaceae;g__Dietzia k__Bacteria;p__Actinobacteria;c__Actinobacteria;o__Actinomycetales;f__Frankiaceae;g__ k__Bacteria;p__Actinobacteria;c__Actinobacteria;o__Actinomycetales;f__Geodermatophilaceae;__ k__Bacteria;p__Actinobacteria;c__Actinobacteria;o__Actinomycetales;f__Geodermatophilaceae;g__ k__Bacteria;p__Actinobacteria;c__Actinobacteria;o__Actinomycetales;f__Geodermatophilaceae;g__Blastococcus k__Bacteria;p__Actinobacteria;c__Actinobacteria;o__Actinomycetales;f__Geodermatophilaceae;g__Modestobacter k__Bacteria;p__Actinobacteria;c__Actinobacteria;o__Actinomycetales;f__Gordoniaceae;g__Gordonia k__Bacteria;p__Actinobacteria;c__Actinobacteria;o__Actinomycetales;f__Intrasporangiaceae;__ k__Bacteria;p__Actinobacteria;c__Actinobacteria;o__Actinomycetales;f__Intrasporangiaceae;g__Arsenicicoccus k__Bacteria;p__Actinobacteria;c__Actinobacteria;o__Actinomycetales;f__Intrasporangiaceae;g__Kytococcus k__Bacteria;p__Actinobacteria;c__Actinobacteria;o__Actinomycetales;f__Jonesiaceae;g__ k__Bacteria;p__Actinobacteria;c__Actinobacteria;o__Actinomycetales;f__Kineosporiaceae;__ k__Bacteria;p__Actinobacteria;c__Actinobacteria;o__Actinomycetales;f__Kineosporiaceae;g__ k__Bacteria;p__Actinobacteria;c__Actinobacteria;o__Actinomycetales;f__Kineosporiaceae;g__Kineococcus k__Bacteria;p__Actinobacteria;c__Actinobacteria;o__Actinomycetales;f__Kineosporiaceae;g__Kineosporia k__Bacteria;p__Actinobacteria;c__Actinobacteria;o__Actinomycetales;f__Microbacteriaceae;__ k__Bacteria;p__Actinobacteria;c__Actinobacteria;o__Actinomycetales;f__Microbacteriaceae;g__Leucobacter k__Bacteria;p__Actinobacteria;c__Actinobacteria;o__Actinomycetales;f__Microbacteriaceae;g__Pseudoclavibacter k__Bacteria;p__Actinobacteria;c__Actinobacteria;o__Actinomycetales;f__Microbacteriaceae;g__Rathayibacter k__Bacteria;p__Actinobacteria;c__Actinobacteria;o__Actinomycetales;f__Micrococcaceae;__ k__Bacteria;p__Actinobacteria;c__Actinobacteria;o__Actinomycetales;f__Micrococcaceae;g__Arthrobacter k__Bacteria;p__Actinobacteria;c__Actinobacteria;o__Actinomycetales;f__Micrococcaceae;g__Kocuria k__Bacteria;p__Actinobacteria;c__Actinobacteria;o__Actinomycetales;f__Micrococcaceae;g__Microbispora k__Bacteria;p__Actinobacteria;c__Actinobacteria;o__Actinomycetales;f__Micrococcaceae;g__Micrococcus k__Bacteria;p__Actinobacteria;c__Actinobacteria;o__Actinomycetales;f__Micrococcaceae;g__Nesterenkonia k__Bacteria;p__Actinobacteria;c__Actinobacteria;o__Actinomycetales;f__Micrococcaceae;g__Rothia k__Bacteria;p__Actinobacteria;c__Actinobacteria;o__Actinomycetales;f__Mycobacteriaceae;g__Mycobacterium k__Bacteria;p__Actinobacteria;c__Actinobacteria;o__Actinomycetales;f__Nakamurellaceae;g__ k__Bacteria;p__Actinobacteria;c__Actinobacteria;o__Actinomycetales;f__Nocardiaceae;g__Rhodococcus k__Bacteria;p__Actinobacteria;c__Actinobacteria;o__Actinomycetales;f__Nocardioidaceae;g__ k__Bacteria;p__Actinobacteria;c__Actinobacteria;o__Actinomycetales;f__Nocardioidaceae;g__Nocardioides k__Bacteria;p__Actinobacteria;c__Actinobacteria;o__Actinomycetales;f__Promicromonosporaceae;g__Promicromonospora k__Bacteria;p__Actinobacteria;c__Actinobacteria;o__Actinomycetales;f__Propionibacteriaceae;g__Propionibacterium k__Bacteria;p__Actinobacteria;c__Actinobacteria;o__Actinomycetales;f__Pseudonocardiaceae;g__Saccharopolyspora k__Bacteria;p__Actinobacteria;c__Actinobacteria;o__Actinomycetales;f__Sporichthyaceae;g__ k__Bacteria;p__Actinobacteria;c__Actinobacteria;o__Actinomycetales;f__Streptomycetaceae;g__Streptomyces k__Bacteria;p__Actinobacteria;c__Actinobacteria;o__Actinomycetales;f__Tsukamurellaceae;g__Tsukamurella k__Bacteria;p__Actinobacteria;c__Actinobacteria;o__Actinomycetales;f__Williamsiaceae;g__Williamsia

k__Bacteria;p__Actinobacteria;c__Actinobacteria;o__Bifidobacteriales;f__Bifidobacteriaceae;g__ k__Bacteria;p__Actinobacteria;c__Actinobacteria;o__Bifidobacteriales;f__Bifidobacteriaceae;g__Alloscardovia k__Bacteria;p__Actinobacteria;c__Actinobacteria;o__Bifidobacteriales;f__Bifidobacteriaceae;g__Bifidobacterium k__Bacteria;p__Actinobacteria;c__Actinobacteria;o__Bifidobacteriales;f__Bifidobacteriaceae;g__Gardnerella k__Bacteria;p__Actinobacteria;c__Actinobacteria;o__Bifidobacteriales;f__Bifidobacteriaceae;g__Scardovia k__Bacteria;p__Actinobacteria;c__Coriobacteriia;o__Coriobacteriales;f__Coriobacteriaceae;__ k__Bacteria;p__Actinobacteria;c__Coriobacteriia;o__Coriobacteriales;f__Coriobacteriaceae;g__ k__Bacteria;p__Actinobacteria;c__Coriobacteriia;o__Coriobacteriales;f__Coriobacteriaceae;g__Atopobium k__Bacteria;p__Actinobacteria;c__Coriobacteriia;o__Coriobacteriales;f__Coriobacteriaceae;g__Collinsella k__Bacteria;p__Actinobacteria;c__Coriobacteriia;o__Coriobacteriales;f__Coriobacteriaceae;g__Olsenella k__Bacteria;p__Actinobacteria;c__Rubrobacteria;o__Rubrobacterales;f__Rubrobacteraceae;g__Rubrobacter k__Bacteria;p__Actinobacteria;c__Thermoleophilia;o__Gaiellales;f__AK1AB1_02E;g__ k__Bacteria;p__Actinobacteria;c__Thermoleophilia;o__Solirubrobacterales;f__Solirubrobacteraceae;g__ k__Bacteria;p__Bacteroidetes;__;__;__;__
k__Bacteria;p__Bacteroidetes;c__Bacteroidia;o__Bacteroidales;__;__
k__Bacteria;p__Bacteroidetes;c__Bacteroidia;o__Bacteroidales;f__;g__
k__Bacteria;p__Bacteroidetes;c__Bacteroidia;o__Bacteroidales;f__Bacteroidaceae;g__Bacteroides k__Bacteria;p__Bacteroidetes;c__Bacteroidia;o__Bacteroidales;f__Porphyromonadaceae;__ k__Bacteria;p__Bacteroidetes;c__Bacteroidia;o__Bacteroidales;f__Porphyromonadaceae;g__Candidatus Azobacteroides k__Bacteria;p__Bacteroidetes;c__Bacteroidia;o__Bacteroidales;f__Porphyromonadaceae;g__Dysgonomonas k__Bacteria;p__Bacteroidetes;c__Bacteroidia;o__Bacteroidales;f__Porphyromonadaceae;g__Paludibacter k__Bacteria;p__Bacteroidetes;c__Bacteroidia;o__Bacteroidales;f__Porphyromonadaceae;g__Parabacteroides k__Bacteria;p__Bacteroidetes;c__Bacteroidia;o__Bacteroidales;f__Porphyromonadaceae;g__Porphyromonas k__Bacteria;p__Bacteroidetes;c__Bacteroidia;o__Bacteroidales;f__Porphyromonadaceae;g__Tannerella k__Bacteria;p__Bacteroidetes;c__Bacteroidia;o__Bacteroidales;f__Prevotellaceae;g__Prevotella k__Bacteria;p__Bacteroidetes;c__Bacteroidia;o__Bacteroidales;f__Rikenellaceae;__ k__Bacteria;p__Bacteroidetes;c__Bacteroidia;o__Bacteroidales;f__Rikenellaceae;g__ k__Bacteria;p__Bacteroidetes;c__Bacteroidia;o__Bacteroidales;f__Rikenellaceae;g__Alistipes k__Bacteria;p__Bacteroidetes;c__Bacteroidia;o__Bacteroidales;f__S24-7;g__
k__Bacteria;p__Bacteroidetes;c__Bacteroidia;o__Bacteroidales;f__[Barnesiellaceae];__ k__Bacteria;p__Bacteroidetes;c__Bacteroidia;o__Bacteroidales;f__[Barnesiellaceae];g__ k__Bacteria;p__Bacteroidetes;c__Bacteroidia;o__Bacteroidales;f__[Odoribacteraceae];g__Odoribacter k__Bacteria;p__Bacteroidetes;c__Bacteroidia;o__Bacteroidales;f__[Paraprevotellaceae];__ k__Bacteria;p__Bacteroidetes;c__Bacteroidia;o__Bacteroidales;f__[Paraprevotellaceae];g__ k__Bacteria;p__Bacteroidetes;c__Bacteroidia;o__Bacteroidales;f__[Paraprevotellaceae];g__[Prevotella] k__Bacteria;p__Bacteroidetes;c__Cytophagia;o__Cytophagales;f__Cytophagaceae;g__Dyadobacter k__Bacteria;p__Bacteroidetes;c__Cytophagia;o__Cytophagales;f__Cytophagaceae;g__Hymenobacter k__Bacteria;p__Bacteroidetes;c__Flavobacteriia;o__Flavobacteriales;__;__
k__Bacteria;p__Bacteroidetes;c__Flavobacteriia;o__Flavobacteriales;f__Blattabacteriaceae;g__Blattabacterium k__Bacteria;p__Bacteroidetes;c__Flavobacteriia;o__Flavobacteriales;f__Flavobacteriaceae;g__Capnocytophaga k__Bacteria;p__Bacteroidetes;c__Flavobacteriia;o__Flavobacteriales;f__Flavobacteriaceae;g__Flavobacterium k__Bacteria;p__Bacteroidetes;c__Flavobacteriia;o__Flavobacteriales;f__Flavobacteriaceae;g__Myroides k__Bacteria;p__Bacteroidetes;c__Flavobacteriia;o__Flavobacteriales;f__[Weeksellaceae];__ k__Bacteria;p__Bacteroidetes;c__Flavobacteriia;o__Flavobacteriales;f__[Weeksellaceae];g__ k__Bacteria;p__Bacteroidetes;c__Flavobacteriia;o__Flavobacteriales;f__[Weeksellaceae];g__Chryseobacterium k__Bacteria;p__Bacteroidetes;c__Flavobacteriia;o__Flavobacteriales;f__[Weeksellaceae];g__Cloacibacterium k__Bacteria;p__Bacteroidetes;c__Flavobacteriia;o__Flavobacteriales;f__[Weeksellaceae];g__Wautersiella k__Bacteria;p__Bacteroidetes;c__Sphingobacteriia;o__Sphingobacteriales;f__;g__ k__Bacteria;p__Bacteroidetes;c__Sphingobacteriia;o__Sphingobacteriales;f__Sphingobacteriaceae;g__Mucilaginibacter k__Bacteria;p__Bacteroidetes;c__Sphingobacteriia;o__Sphingobacteriales;f__Sphingobacteriaceae;g__Pedobacter k__Bacteria;p__Bacteroidetes;c__Sphingobacteriia;o__Sphingobacteriales;f__Sphingobacteriaceae;g__Sphingobacterium k__Bacteria;p__Bacteroidetes;c__[Saprospirae];o__[Saprospirales];f__Chitinophagaceae;__ k__Bacteria;p__Bacteroidetes;c__[Saprospirae];o__[Saprospirales];f__Chitinophagaceae;g__Flavisolibacter k__Bacteria;p__Bacteroidetes;c__[Saprospirae];o__[Saprospirales];f__Chitinophagaceae;g__Sediminibacterium k__Bacteria;p__Bacteroidetes;c__[Saprospirae];o__[Saprospirales];f__Chitinophagaceae;g__Segetibacter k__Bacteria;p__Chlamydiae;c__Chlamydiia;o__Chlamydiales;f__Parachlamydiaceae;g__Candidatus Protochlamydia k__Bacteria;p__Chloroflexi;c__Anaerolineae;o__Anaerolineales;f__Anaerolinaceae;g__SHD-231 k__Bacteria;p__Chloroflexi;c__Gitt-GS-136;o__;f__;g__
k__Bacteria;p__Chloroflexi;c__Thermomicrobia;o__JG30-KF-CM45;f__;g__
k__Bacteria;p__Cyanobacteria;c__4C0d-2;o__MLE1-12;f__;g__
k__Bacteria;p__Cyanobacteria;c__4C0d-2;o__YS2;f__;g__

k__Bacteria;p__Cyanobacteria;c__Chloroplast;o__Chlorophyta;f__;g__
k__Bacteria;p__Cyanobacteria;c__Chloroplast;o__Chlorophyta;f__Trebouxiophyceae;g__Chloroidium k__Bacteria;p__Cyanobacteria;c__Chloroplast;o__Haptophyceae;f__;g__
k__Bacteria;p__Cyanobacteria;c__Chloroplast;o__Streptophyta;f__;g__
k__Bacteria;p__Cyanobacteria;c__ML635J-21;o__;f__;g__
k__Bacteria;p__Cyanobacteria;c__Nostocophycideae;o__Nostocales;f__Nostocaceae;__ k__Bacteria;p__Cyanobacteria;c__Oscillatoriophycideae;o__Chroococcales;f__;g__ k__Bacteria;p__Cyanobacteria;c__Oscillatoriophycideae;o__Chroococcales;f__Xenococcaceae;g__Chroococcidiopsis k__Bacteria;p__Cyanobacteria;c__Oscillatoriophycideae;o__Oscillatoriales;f__Phormidiaceae;g__Phormidium k__Bacteria;p__FBP;c__;o__;f__;g__
k__Bacteria;p__Firmicutes;__;__;__;__
k__Bacteria;p__Firmicutes;c__Bacilli;__;__;__
k__Bacteria;p__Firmicutes;c__Bacilli;o__Bacillales;__;__
k__Bacteria;p__Firmicutes;c__Bacilli;o__Bacillales;f__;g__
k__Bacteria;p__Firmicutes;c__Bacilli;o__Bacillales;f__Alicyclobacillaceae;g__Alicyclobacillus k__Bacteria;p__Firmicutes;c__Bacilli;o__Bacillales;f__Bacillaceae;__
k__Bacteria;p__Firmicutes;c__Bacilli;o__Bacillales;f__Bacillaceae;g__Anoxybacillus k__Bacteria;p__Firmicutes;c__Bacilli;o__Bacillales;f__Bacillaceae;g__Bacillus k__Bacteria;p__Firmicutes;c__Bacilli;o__Bacillales;f__Bacillaceae;g__Geobacillus k__Bacteria;p__Firmicutes;c__Bacilli;o__Bacillales;f__Bacillaceae;g__Paucisalibacillus k__Bacteria;p__Firmicutes;c__Bacilli;o__Bacillales;f__Listeriaceae;g__Brochothrix k__Bacteria;p__Firmicutes;c__Bacilli;o__Bacillales;f__Paenibacillaceae;g__Paenibacillus k__Bacteria;p__Firmicutes;c__Bacilli;o__Bacillales;f__Planococcaceae;__
k__Bacteria;p__Firmicutes;c__Bacilli;o__Bacillales;f__Planococcaceae;g__
k__Bacteria;p__Firmicutes;c__Bacilli;o__Bacillales;f__Planococcaceae;g__Bacillus k__Bacteria;p__Firmicutes;c__Bacilli;o__Bacillales;f__Planococcaceae;g__Sporosarcina k__Bacteria;p__Firmicutes;c__Bacilli;o__Bacillales;f__Staphylococcaceae;g__Jeotgalicoccus k__Bacteria;p__Firmicutes;c__Bacilli;o__Bacillales;f__Staphylococcaceae;g__Macrococcus k__Bacteria;p__Firmicutes;c__Bacilli;o__Bacillales;f__Staphylococcaceae;g__Salinicoccus k__Bacteria;p__Firmicutes;c__Bacilli;o__Bacillales;f__Staphylococcaceae;g__Staphylococcus k__Bacteria;p__Firmicutes;c__Bacilli;o__Bacillales;f__Thermoactinomycetaceae;g__Thermoactinomyces k__Bacteria;p__Firmicutes;c__Bacilli;o__Bacillales;f__[Exiguobacteraceae];__ k__Bacteria;p__Firmicutes;c__Bacilli;o__Bacillales;f__[Exiguobacteraceae];g__Exiguobacterium k__Bacteria;p__Firmicutes;c__Bacilli;o__Gemellales;f__Gemellaceae;__
k__Bacteria;p__Firmicutes;c__Bacilli;o__Gemellales;f__Gemellaceae;g__
k__Bacteria;p__Firmicutes;c__Bacilli;o__Lactobacillales;__;__
k__Bacteria;p__Firmicutes;c__Bacilli;o__Lactobacillales;f__Aerococcaceae;__
k__Bacteria;p__Firmicutes;c__Bacilli;o__Lactobacillales;f__Aerococcaceae;g__Abiotrophia k__Bacteria;p__Firmicutes;c__Bacilli;o__Lactobacillales;f__Aerococcaceae;g__Aerococcus k__Bacteria;p__Firmicutes;c__Bacilli;o__Lactobacillales;f__Aerococcaceae;g__Alloiococcus k__Bacteria;p__Firmicutes;c__Bacilli;o__Lactobacillales;f__Aerococcaceae;g__Facklamia k__Bacteria;p__Firmicutes;c__Bacilli;o__Lactobacillales;f__Carnobacteriaceae;g__Granulicatella k__Bacteria;p__Firmicutes;c__Bacilli;o__Lactobacillales;f__Enterococcaceae;g__Enterococcus k__Bacteria;p__Firmicutes;c__Bacilli;o__Lactobacillales;f__Enterococcaceae;g__Tetragenococcus k__Bacteria;p__Firmicutes;c__Bacilli;o__Lactobacillales;f__Lactobacillaceae;__ k__Bacteria;p__Firmicutes;c__Bacilli;o__Lactobacillales;f__Lactobacillaceae;g__Lactobacillus k__Bacteria;p__Firmicutes;c__Bacilli;o__Lactobacillales;f__Lactobacillaceae;g__Pediococcus k__Bacteria;p__Firmicutes;c__Bacilli;o__Lactobacillales;f__Leuconostocaceae;__ k__Bacteria;p__Firmicutes;c__Bacilli;o__Lactobacillales;f__Leuconostocaceae;g__Leuconostoc k__Bacteria;p__Firmicutes;c__Bacilli;o__Lactobacillales;f__Leuconostocaceae;g__Weissella k__Bacteria;p__Firmicutes;c__Bacilli;o__Lactobacillales;f__Streptococcaceae;__ k__Bacteria;p__Firmicutes;c__Bacilli;o__Lactobacillales;f__Streptococcaceae;g__ k__Bacteria;p__Firmicutes;c__Bacilli;o__Lactobacillales;f__Streptococcaceae;g__Lactococcus k__Bacteria;p__Firmicutes;c__Bacilli;o__Lactobacillales;f__Streptococcaceae;g__Streptococcus k__Bacteria;p__Firmicutes;c__Bacilli;o__Turicibacterales;f__Turicibacteraceae;g__Turicibacter k__Bacteria;p__Firmicutes;c__Clostridia;__;__;__
k__Bacteria;p__Firmicutes;c__Clostridia;o__Clostridiales;__;__
k__Bacteria;p__Firmicutes;c__Clostridia;o__Clostridiales;f__;g__
k__Bacteria;p__Firmicutes;c__Clostridia;o__Clostridiales;f__Christensenellaceae;g__ k__Bacteria;p__Firmicutes;c__Clostridia;o__Clostridiales;f__Clostridiaceae;__ k__Bacteria;p__Firmicutes;c__Clostridia;o__Clostridiales;f__Clostridiaceae;g__

k__Bacteria;p__Firmicutes;c__Clostridia;o__Clostridiales;f__Clostridiaceae;g__Caloramator k__Bacteria;p__Firmicutes;c__Clostridia;o__Clostridiales;f__Clostridiaceae;g__Clostridium k__Bacteria;p__Firmicutes;c__Clostridia;o__Clostridiales;f__Clostridiaceae;g__Thermoanaerobacterium k__Bacteria;p__Firmicutes;c__Clostridia;o__Clostridiales;f__Eubacteriaceae;g__Pseudoramibacter_Eubacterium k__Bacteria;p__Firmicutes;c__Clostridia;o__Clostridiales;f__Lachnospiraceae;__ k__Bacteria;p__Firmicutes;c__Clostridia;o__Clostridiales;f__Lachnospiraceae;g__ k__Bacteria;p__Firmicutes;c__Clostridia;o__Clostridiales;f__Lachnospiraceae;g__Anaerostipes k__Bacteria;p__Firmicutes;c__Clostridia;o__Clostridiales;f__Lachnospiraceae;g__Blautia k__Bacteria;p__Firmicutes;c__Clostridia;o__Clostridiales;f__Lachnospiraceae;g__Butyrivibrio k__Bacteria;p__Firmicutes;c__Clostridia;o__Clostridiales;f__Lachnospiraceae;g__Clostridium k__Bacteria;p__Firmicutes;c__Clostridia;o__Clostridiales;f__Lachnospiraceae;g__Coprococcus k__Bacteria;p__Firmicutes;c__Clostridia;o__Clostridiales;f__Lachnospiraceae;g__Dorea k__Bacteria;p__Firmicutes;c__Clostridia;o__Clostridiales;f__Lachnospiraceae;g__Johnsonella k__Bacteria;p__Firmicutes;c__Clostridia;o__Clostridiales;f__Lachnospiraceae;g__Lachnoanaerobaculum k__Bacteria;p__Firmicutes;c__Clostridia;o__Clostridiales;f__Lachnospiraceae;g__Lachnobacterium k__Bacteria;p__Firmicutes;c__Clostridia;o__Clostridiales;f__Lachnospiraceae;g__Lachnospira k__Bacteria;p__Firmicutes;c__Clostridia;o__Clostridiales;f__Lachnospiraceae;g__Moryella k__Bacteria;p__Firmicutes;c__Clostridia;o__Clostridiales;f__Lachnospiraceae;g__Oribacterium k__Bacteria;p__Firmicutes;c__Clostridia;o__Clostridiales;f__Lachnospiraceae;g__Roseburia k__Bacteria;p__Firmicutes;c__Clostridia;o__Clostridiales;f__Lachnospiraceae;g__Ruminococcus k__Bacteria;p__Firmicutes;c__Clostridia;o__Clostridiales;f__Lachnospiraceae;g__Shuttleworthia k__Bacteria;p__Firmicutes;c__Clostridia;o__Clostridiales;f__Lachnospiraceae;g__[Ruminococcus] k__Bacteria;p__Firmicutes;c__Clostridia;o__Clostridiales;f__Peptococcaceae;g__Peptococcus k__Bacteria;p__Firmicutes;c__Clostridia;o__Clostridiales;f__Peptostreptococcaceae;__ k__Bacteria;p__Firmicutes;c__Clostridia;o__Clostridiales;f__Peptostreptococcaceae;g__ k__Bacteria;p__Firmicutes;c__Clostridia;o__Clostridiales;f__Peptostreptococcaceae;g__Clostridium k__Bacteria;p__Firmicutes;c__Clostridia;o__Clostridiales;f__Peptostreptococcaceae;g__Filifactor k__Bacteria;p__Firmicutes;c__Clostridia;o__Clostridiales;f__Peptostreptococcaceae;g__Peptostreptococcus k__Bacteria;p__Firmicutes;c__Clostridia;o__Clostridiales;f__Peptostreptococcaceae;g__[Clostridium] k__Bacteria;p__Firmicutes;c__Clostridia;o__Clostridiales;f__Ruminococcaceae;__ k__Bacteria;p__Firmicutes;c__Clostridia;o__Clostridiales;f__Ruminococcaceae;g__ k__Bacteria;p__Firmicutes;c__Clostridia;o__Clostridiales;f__Ruminococcaceae;g__Clostridium k__Bacteria;p__Firmicutes;c__Clostridia;o__Clostridiales;f__Ruminococcaceae;g__Faecalibacterium k__Bacteria;p__Firmicutes;c__Clostridia;o__Clostridiales;f__Ruminococcaceae;g__Gemmiger k__Bacteria;p__Firmicutes;c__Clostridia;o__Clostridiales;f__Ruminococcaceae;g__Oscillospira k__Bacteria;p__Firmicutes;c__Clostridia;o__Clostridiales;f__Ruminococcaceae;g__Ruminococcus k__Bacteria;p__Firmicutes;c__Clostridia;o__Clostridiales;f__Veillonellaceae;__ k__Bacteria;p__Firmicutes;c__Clostridia;o__Clostridiales;f__Veillonellaceae;g__ k__Bacteria;p__Firmicutes;c__Clostridia;o__Clostridiales;f__Veillonellaceae;g__Acidaminococcus k__Bacteria;p__Firmicutes;c__Clostridia;o__Clostridiales;f__Veillonellaceae;g__Anaerovibrio k__Bacteria;p__Firmicutes;c__Clostridia;o__Clostridiales;f__Veillonellaceae;g__Dialister k__Bacteria;p__Firmicutes;c__Clostridia;o__Clostridiales;f__Veillonellaceae;g__Megasphaera k__Bacteria;p__Firmicutes;c__Clostridia;o__Clostridiales;f__Veillonellaceae;g__Phascolarctobacterium k__Bacteria;p__Firmicutes;c__Clostridia;o__Clostridiales;f__Veillonellaceae;g__Schwartzia k__Bacteria;p__Firmicutes;c__Clostridia;o__Clostridiales;f__Veillonellaceae;g__Selenomonas k__Bacteria;p__Firmicutes;c__Clostridia;o__Clostridiales;f__Veillonellaceae;g__Veillonella k__Bacteria;p__Firmicutes;c__Clostridia;o__Clostridiales;f__[Mogibacteriaceae];__ k__Bacteria;p__Firmicutes;c__Clostridia;o__Clostridiales;f__[Mogibacteriaceae];g__ k__Bacteria;p__Firmicutes;c__Clostridia;o__Clostridiales;f__[Mogibacteriaceae];g__Anaerovorax k__Bacteria;p__Firmicutes;c__Clostridia;o__Clostridiales;f__[Mogibacteriaceae];g__Mogibacterium k__Bacteria;p__Firmicutes;c__Clostridia;o__Clostridiales;f__[Tissierellaceae];g__ k__Bacteria;p__Firmicutes;c__Clostridia;o__Clostridiales;f__[Tissierellaceae];g__1-68 k__Bacteria;p__Firmicutes;c__Clostridia;o__Clostridiales;f__[Tissierellaceae];g__Anaerococcus k__Bacteria;p__Firmicutes;c__Clostridia;o__Clostridiales;f__[Tissierellaceae];g__Finegoldia k__Bacteria;p__Firmicutes;c__Clostridia;o__Clostridiales;f__[Tissierellaceae];g__Gallicola k__Bacteria;p__Firmicutes;c__Clostridia;o__Clostridiales;f__[Tissierellaceae];g__Parvimonas k__Bacteria;p__Firmicutes;c__Clostridia;o__Clostridiales;f__[Tissierellaceae];g__Peptoniphilus k__Bacteria;p__Firmicutes;c__Clostridia;o__Clostridiales;f__[Tissierellaceae];g__WAL_1855D k__Bacteria;p__Firmicutes;c__Clostridia;o__Clostridiales;f__[Tissierellaceae];g__ph2 k__Bacteria;p__Firmicutes;c__Clostridia;o__Thermoanaerobacterales;f__Caldicellulosiruptoraceae;g__Caldicellulosiruptor k__Bacteria;p__Firmicutes;c__Clostridia;o__Thermoanaerobacterales;f__Thermoanaerobacteraceae;g__Caldanaerobacter

k__Bacteria;p__Firmicutes;c__Erysipelotrichi;o__Erysipelotrichales;f__Erysipelotrichaceae;g__ k__Bacteria;p__Firmicutes;c__Erysipelotrichi;o__Erysipelotrichales;f__Erysipelotrichaceae;g__Bulleidia k__Bacteria;p__Firmicutes;c__Erysipelotrichi;o__Erysipelotrichales;f__Erysipelotrichaceae;g__Catenibacterium k__Bacteria;p__Firmicutes;c__Erysipelotrichi;o__Erysipelotrichales;f__Erysipelotrichaceae;g__Clostridium k__Bacteria;p__Firmicutes;c__Erysipelotrichi;o__Erysipelotrichales;f__Erysipelotrichaceae;g__Erysipelothrix k__Bacteria;p__Firmicutes;c__Erysipelotrichi;o__Erysipelotrichales;f__Erysipelotrichaceae;g__Sharpea k__Bacteria;p__Firmicutes;c__Erysipelotrichi;o__Erysipelotrichales;f__Erysipelotrichaceae;g__[Eubacterium] k__Bacteria;p__Firmicutes;c__Erysipelotrichi;o__Erysipelotrichales;f__Erysipelotrichaceae;g__cc_115 k__Bacteria;p__Fusobacteria;c__Fusobacteriia;o__Fusobacteriales;__;__
k__Bacteria;p__Fusobacteria;c__Fusobacteriia;o__Fusobacteriales;f__Fusobacteriaceae;g__Fusobacterium k__Bacteria;p__Fusobacteria;c__Fusobacteriia;o__Fusobacteriales;f__Leptotrichiaceae;__ k__Bacteria;p__Fusobacteria;c__Fusobacteriia;o__Fusobacteriales;f__Leptotrichiaceae;g__ k__Bacteria;p__Fusobacteria;c__Fusobacteriia;o__Fusobacteriales;f__Leptotrichiaceae;g__Leptotrichia k__Bacteria;p__Fusobacteria;c__Fusobacteriia;o__Fusobacteriales;f__Leptotrichiaceae;g__Sneathia k__Bacteria;p__GN02;c__BD1-5;o__;f__;g__
k__Bacteria;p__Gemmatimonadetes;c__Gemmatimonadetes;__;__;__
k__Bacteria;p__OD1;c__;o__;f__;g__
k__Bacteria;p__Planctomycetes;c__Phycisphaerae;o__WD2101;f__;g__
k__Bacteria;p__Proteobacteria;__;__;__;__
k__Bacteria;p__Proteobacteria;c__Alphaproteobacteria;__;__;__
k__Bacteria;p__Proteobacteria;c__Alphaproteobacteria;o__Caulobacterales;f__Caulobacteraceae;g__Brevundimonas k__Bacteria;p__Proteobacteria;c__Alphaproteobacteria;o__Caulobacterales;f__Caulobacteraceae;g__Caulobacter k__Bacteria;p__Proteobacteria;c__Alphaproteobacteria;o__Caulobacterales;f__Caulobacteraceae;g__Mycoplana k__Bacteria;p__Proteobacteria;c__Alphaproteobacteria;o__Caulobacterales;f__Caulobacteraceae;g__Phenylobacterium k__Bacteria;p__Proteobacteria;c__Alphaproteobacteria;o__RF32;f__;g__
k__Bacteria;p__Proteobacteria;c__Alphaproteobacteria;o__Rhodobacterales;f__Rhodobacteraceae;__ k__Bacteria;p__Proteobacteria;c__Alphaproteobacteria;o__Rhodobacterales;f__Rhodobacteraceae;g__Paracoccus k__Bacteria;p__Proteobacteria;c__Alphaproteobacteria;o__Rhodobacterales;f__Rhodobacteraceae;g__Rhodobacter k__Bacteria;p__Proteobacteria;c__Alphaproteobacteria;o__Rhodobacterales;f__Rhodobacteraceae;g__Rubellimicrobium k__Bacteria;p__Proteobacteria;c__Alphaproteobacteria;o__Rhodospirillales;f__;g__ k__Bacteria;p__Proteobacteria;c__Alphaproteobacteria;o__Rhodospirillales;f__Acetobacteraceae;__ k__Bacteria;p__Proteobacteria;c__Alphaproteobacteria;o__Rhodospirillales;f__Acetobacteraceae;g__ k__Bacteria;p__Proteobacteria;c__Alphaproteobacteria;o__Rhodospirillales;f__Acetobacteraceae;g__Acetobacter k__Bacteria;p__Proteobacteria;c__Alphaproteobacteria;o__Rhodospirillales;f__Acetobacteraceae;g__Gluconacetobacter k__Bacteria;p__Proteobacteria;c__Alphaproteobacteria;o__Rhodospirillales;f__Acetobacteraceae;g__Gluconobacter k__Bacteria;p__Proteobacteria;c__Alphaproteobacteria;o__Rhodospirillales;f__Acetobacteraceae;g__Roseomonas k__Bacteria;p__Proteobacteria;c__Alphaproteobacteria;o__Rhodospirillales;f__Acetobacteraceae;g__Swaminathania k__Bacteria;p__Proteobacteria;c__Alphaproteobacteria;o__Rhodospirillales;f__Acetobacteraceae;g__Tanticharoenia k__Bacteria;p__Proteobacteria;c__Alphaproteobacteria;o__Rhodospirillales;f__Rhodospirillaceae;g__Azospirillum k__Bacteria;p__Proteobacteria;c__Alphaproteobacteria;o__Rhodospirillales;f__Rhodospirillaceae;g__Reyranella k__Bacteria;p__Proteobacteria;c__Alphaproteobacteria;o__Rhodospirillales;f__Rhodospirillaceae;g__Skermanella k__Bacteria;p__Proteobacteria;c__Alphaproteobacteria;o__Rickettsiales;f__;g__ k__Bacteria;p__Proteobacteria;c__Alphaproteobacteria;o__Rickettsiales;f__Rickettsiaceae;g__Wolbachia k__Bacteria;p__Proteobacteria;c__Alphaproteobacteria;o__Rickettsiales;f__mitochondria;__ k__Bacteria;p__Proteobacteria;c__Alphaproteobacteria;o__Rickettsiales;f__mitochondria;g__ k__Bacteria;p__Proteobacteria;c__Alphaproteobacteria;o__Rickettsiales;f__mitochondria;g__Acanthamoeba k__Bacteria;p__Proteobacteria;c__Alphaproteobacteria;o__Rickettsiales;f__mitochondria;g__Malus k__Bacteria;p__Proteobacteria;c__Alphaproteobacteria;o__Rickettsiales;f__mitochondria;g__Phytophthora k__Bacteria;p__Proteobacteria;c__Alphaproteobacteria;o__Sphingomonadales;__;__ k__Bacteria;p__Proteobacteria;c__Alphaproteobacteria;o__Sphingomonadales;f__Erythrobacteraceae;__ k__Bacteria;p__Proteobacteria;c__Alphaproteobacteria;o__Sphingomonadales;f__Erythrobacteraceae;g__Erythrobacter k__Bacteria;p__Proteobacteria;c__Alphaproteobacteria;o__Sphingomonadales;f__Erythrobacteraceae;g__Lutibacterium k__Bacteria;p__Proteobacteria;c__Alphaproteobacteria;o__Sphingomonadales;f__Sphingomonadaceae;__ k__Bacteria;p__Proteobacteria;c__Alphaproteobacteria;o__Sphingomonadales;f__Sphingomonadaceae;g__Kaistobacter k__Bacteria;p__Proteobacteria;c__Alphaproteobacteria;o__Sphingomonadales;f__Sphingomonadaceae;g__Novosphingobium k__Bacteria;p__Proteobacteria;c__Alphaproteobacteria;o__Sphingomonadales;f__Sphingomonadaceae;g__Sandaracinobacter k__Bacteria;p__Proteobacteria;c__Alphaproteobacteria;o__Sphingomonadales;f__Sphingomonadaceae;g__Sphingobium k__Bacteria;p__Proteobacteria;c__Alphaproteobacteria;o__Sphingomonadales;f__Sphingomonadaceae;g__Sphingomonas k__Bacteria;p__Proteobacteria;c__Betaproteobacteria;__;__;__
k__Bacteria;p__Proteobacteria;c__Betaproteobacteria;o__;f__;g__
k__Bacteria;p__Proteobacteria;c__Betaproteobacteria;o__Burkholderiales;f__Alcaligenaceae;__

k__Bacteria;p__Proteobacteria;c__Betaproteobacteria;o__Burkholderiales;f__Alcaligenaceae;g__Achromobacter k__Bacteria;p__Proteobacteria;c__Betaproteobacteria;o__Burkholderiales;f__Alcaligenaceae;g__Sutterella k__Bacteria;p__Proteobacteria;c__Betaproteobacteria;o__Burkholderiales;f__Burkholderiaceae;g__Burkholderia k__Bacteria;p__Proteobacteria;c__Betaproteobacteria;o__Burkholderiales;f__Burkholderiaceae;g__Lautropia k__Bacteria;p__Proteobacteria;c__Betaproteobacteria;o__Burkholderiales;f__Comamonadaceae;__ k__Bacteria;p__Proteobacteria;c__Betaproteobacteria;o__Burkholderiales;f__Comamonadaceae;g__Comamonas k__Bacteria;p__Proteobacteria;c__Betaproteobacteria;o__Burkholderiales;f__Comamonadaceae;g__Tepidimonas k__Bacteria;p__Proteobacteria;c__Betaproteobacteria;o__Burkholderiales;f__Oxalobacteraceae;__ k__Bacteria;p__Proteobacteria;c__Betaproteobacteria;o__Burkholderiales;f__Oxalobacteraceae;g__Herbaspirillum k__Bacteria;p__Proteobacteria;c__Betaproteobacteria;o__Burkholderiales;f__Oxalobacteraceae;g__Massilia k__Bacteria;p__Proteobacteria;c__Betaproteobacteria;o__Methylophilales;f__Methylophilaceae;g__Methylotenera k__Bacteria;p__Proteobacteria;c__Betaproteobacteria;o__Neisseriales;f__Neisseriaceae;__ k__Bacteria;p__Proteobacteria;c__Betaproteobacteria;o__Neisseriales;f__Neisseriaceae;g__ k__Bacteria;p__Proteobacteria;c__Betaproteobacteria;o__Neisseriales;f__Neisseriaceae;g__Conchiformibius k__Bacteria;p__Proteobacteria;c__Betaproteobacteria;o__Neisseriales;f__Neisseriaceae;g__Eikenella k__Bacteria;p__Proteobacteria;c__Betaproteobacteria;o__Neisseriales;f__Neisseriaceae;g__Gulbenkiania k__Bacteria;p__Proteobacteria;c__Betaproteobacteria;o__Neisseriales;f__Neisseriaceae;g__Kingella k__Bacteria;p__Proteobacteria;c__Betaproteobacteria;o__Neisseriales;f__Neisseriaceae;g__Neisseria k__Bacteria;p__Proteobacteria;c__Betaproteobacteria;o__Rhodocyclales;f__Rhodocyclaceae;__ k__Bacteria;p__Proteobacteria;c__Betaproteobacteria;o__Rhodocyclales;f__Rhodocyclaceae;g__Azospira k__Bacteria;p__Proteobacteria;c__Betaproteobacteria;o__Rhodocyclales;f__Rhodocyclaceae;g__Zoogloea k__Bacteria;p__Proteobacteria;c__Deltaproteobacteria;o__Bdellovibrionales;f__Bacteriovoracaceae;__ k__Bacteria;p__Proteobacteria;c__Deltaproteobacteria;o__Bdellovibrionales;f__Bdellovibrionaceae;g__Bdellovibrio k__Bacteria;p__Proteobacteria;c__Deltaproteobacteria;o__Desulfobacterales;f__Desulfobulbaceae;g__Desulfobulbus k__Bacteria;p__Proteobacteria;c__Deltaproteobacteria;o__Desulfovibrionales;f__Desulfomicrobiaceae;g__ k__Bacteria;p__Proteobacteria;c__Deltaproteobacteria;o__Desulfovibrionales;f__Desulfomicrobiaceae;g__Desulfomicrobium k__Bacteria;p__Proteobacteria;c__Deltaproteobacteria;o__Desulfovibrionales;f__Desulfovibrionaceae;__ k__Bacteria;p__Proteobacteria;c__Deltaproteobacteria;o__Desulfovibrionales;f__Desulfovibrionaceae;g__Desulfovibrio k__Bacteria;p__Proteobacteria;c__Deltaproteobacteria;o__GMD14H09;f__;g__
k__Bacteria;p__Proteobacteria;c__Deltaproteobacteria;o__MIZ46;f__;g__
k__Bacteria;p__Proteobacteria;c__Deltaproteobacteria;o__Myxococcales;__;__
k__Bacteria;p__Proteobacteria;c__Deltaproteobacteria;o__Myxococcales;f__0319-6G20;g__ k__Bacteria;p__Proteobacteria;c__Deltaproteobacteria;o__Myxococcales;f__;g__ k__Bacteria;p__Proteobacteria;c__Deltaproteobacteria;o__Thermodesulfobacteriales;f__Thermodesulfobacteriaceae;g__ k__Bacteria;p__Proteobacteria;c__Epsilonproteobacteria;o__Campylobacterales;f__Campylobacteraceae;g__Campylobacter k__Bacteria;p__Proteobacteria;c__Gammaproteobacteria;__;__;__
k__Bacteria;p__Proteobacteria;c__Gammaproteobacteria;o__Aeromonadales;f__Succinivibrionaceae;g__Succinivibrio k__Bacteria;p__Proteobacteria;c__Gammaproteobacteria;o__Alteromonadales;f__Psychromonadaceae;g__Psychromonas k__Bacteria;p__Proteobacteria;c__Gammaproteobacteria;o__Alteromonadales;f__Shewanellaceae;g__Shewanella k__Bacteria;p__Proteobacteria;c__Gammaproteobacteria;o__Alteromonadales;f__[Chromatiaceae];__ k__Bacteria;p__Proteobacteria;c__Gammaproteobacteria;o__Alteromonadales;f__[Chromatiaceae];g__Alishewanella k__Bacteria;p__Proteobacteria;c__Gammaproteobacteria;o__Cardiobacteriales;f__Cardiobacteriaceae;g__ k__Bacteria;p__Proteobacteria;c__Gammaproteobacteria;o__Cardiobacteriales;f__Cardiobacteriaceae;g__Cardiobacterium k__Bacteria;p__Proteobacteria;c__Gammaproteobacteria;o__Chromatiales;f__Ectothiorhodospiraceae;g__Halorhodospira k__Bacteria;p__Proteobacteria;c__Gammaproteobacteria;o__Enterobacteriales;f__Enterobacteriaceae;__ k__Bacteria;p__Proteobacteria;c__Gammaproteobacteria;o__Enterobacteriales;f__Enterobacteriaceae;g__Erwinia k__Bacteria;p__Proteobacteria;c__Gammaproteobacteria;o__Enterobacteriales;f__Enterobacteriaceae;g__Proteus k__Bacteria;p__Proteobacteria;c__Gammaproteobacteria;o__Legionellales;f__Coxiellaceae;g__ k__Bacteria;p__Proteobacteria;c__Gammaproteobacteria;o__Legionellales;f__Legionellaceae;__ k__Bacteria;p__Proteobacteria;c__Gammaproteobacteria;o__Legionellales;f__Legionellaceae;g__Legionella k__Bacteria;p__Proteobacteria;c__Gammaproteobacteria;o__Oceanospirillales;f__Halomonadaceae;g__Halomonas k__Bacteria;p__Proteobacteria;c__Gammaproteobacteria;o__Oceanospirillales;f__Oceanospirillaceae;g__Marinomonas k__Bacteria;p__Proteobacteria;c__Gammaproteobacteria;o__Oceanospirillales;f__Oceanospirillaceae;g__Nitrincola k__Bacteria;p__Proteobacteria;c__Gammaproteobacteria;o__Pasteurellales;f__Pasteurellaceae;__ k__Bacteria;p__Proteobacteria;c__Gammaproteobacteria;o__Pasteurellales;f__Pasteurellaceae;g__ k__Bacteria;p__Proteobacteria;c__Gammaproteobacteria;o__Pasteurellales;f__Pasteurellaceae;g__Actinobacillus k__Bacteria;p__Proteobacteria;c__Gammaproteobacteria;o__Pasteurellales;f__Pasteurellaceae;g__Aggregatibacter k__Bacteria;p__Proteobacteria;c__Gammaproteobacteria;o__Pasteurellales;f__Pasteurellaceae;g__Haemophilus k__Bacteria;p__Proteobacteria;c__Gammaproteobacteria;o__Pasteurellales;f__Pasteurellaceae;g__Pasteurella k__Bacteria;p__Proteobacteria;c__Gammaproteobacteria;o__Pseudomonadales;f__Moraxellaceae;__ k__Bacteria;p__Proteobacteria;c__Gammaproteobacteria;o__Pseudomonadales;f__Moraxellaceae;g__

k__Bacteria;p__Proteobacteria;c__Gammaproteobacteria;o__Pseudomonadales;f__Moraxellaceae;g__Acinetobacter k__Bacteria;p__Proteobacteria;c__Gammaproteobacteria;o__Pseudomonadales;f__Moraxellaceae;g__Alkanindiges k__Bacteria;p__Proteobacteria;c__Gammaproteobacteria;o__Pseudomonadales;f__Moraxellaceae;g__Enhydrobacter k__Bacteria;p__Proteobacteria;c__Gammaproteobacteria;o__Pseudomonadales;f__Moraxellaceae;g__Moraxella k__Bacteria;p__Proteobacteria;c__Gammaproteobacteria;o__Pseudomonadales;f__Moraxellaceae;g__Psychrobacter k__Bacteria;p__Proteobacteria;c__Gammaproteobacteria;o__Pseudomonadales;f__Pseudomonadaceae;__ k__Bacteria;p__Proteobacteria;c__Gammaproteobacteria;o__Pseudomonadales;f__Pseudomonadaceae;g__Pseudomonas k__Bacteria;p__Proteobacteria;c__Gammaproteobacteria;o__Vibrionales;f__Pseudoalteromonadaceae;g__Pseudoalteromonas k__Bacteria;p__Proteobacteria;c__Gammaproteobacteria;o__Vibrionales;f__Vibrionaceae;g__Photobacterium k__Bacteria;p__Proteobacteria;c__Gammaproteobacteria;o__Vibrionales;f__Vibrionaceae;g__Vibrio k__Bacteria;p__Proteobacteria;c__Gammaproteobacteria;o__Xanthomonadales;f__Sinobacteraceae;g__ k__Bacteria;p__Proteobacteria;c__Gammaproteobacteria;o__Xanthomonadales;f__Sinobacteraceae;g__Solimonas k__Bacteria;p__Proteobacteria;c__Gammaproteobacteria;o__Xanthomonadales;f__Xanthomonadaceae;__ k__Bacteria;p__Proteobacteria;c__Gammaproteobacteria;o__Xanthomonadales;f__Xanthomonadaceae;g__Pseudoxanthomonas k__Bacteria;p__Proteobacteria;c__Gammaproteobacteria;o__Xanthomonadales;f__Xanthomonadaceae;g__Stenotrophomonas k__Bacteria;p__Proteobacteria;c__Gammaproteobacteria;o__Xanthomonadales;f__Xanthomonadaceae;g__Thermomonas k__Bacteria;p__Proteobacteria;c__Gammaproteobacteria;o__Xanthomonadales;f__Xanthomonadaceae;g__Xanthomonas k__Bacteria;p__SBR1093;c__VHS-B5-50;o__;f__;g__
k__Bacteria;p__SR1;c__;o__;f__;g__
k__Bacteria;p__Spirochaetes;c__Spirochaetes;o__Spirochaetales;f__Spirochaetaceae;g__Treponema k__Bacteria;p__Synergistetes;c__Synergistia;o__Synergistales;f__Dethiosulfovibrionaceae;g__TG5 k__Bacteria;p__TM7;c__TM7-3;o__;f__;g__
k__Bacteria;p__TM7;c__TM7-3;o__CW040;f__F16;g__
k__Bacteria;p__Tenericutes;c__Mollicutes;o__Acholeplasmatales;f__Acholeplasmataceae;g__Acholeplasma k__Bacteria;p__Tenericutes;c__Mollicutes;o__Mycoplasmatales;f__Mycoplasmataceae;g__Mycoplasma k__Bacteria;p__Tenericutes;c__Mollicutes;o__RF39;f__;g__
k__Bacteria;p__Tenericutes;c__RF3;o__ML615J-28;f__;g__
k__Bacteria;p__Verrucomicrobia;c__Opitutae;o__Opitutales;f__Opitutaceae;g__Opitutus k__Bacteria;p__Verrucomicrobia;c__Verrucomicrobiae;o__Verrucomicrobiales;f__Verrucomicrobiaceae;g__ k__Bacteria;p__Verrucomicrobia;c__Verrucomicrobiae;o__Verrucomicrobiales;f__Verrucomicrobiaceae;g__Akkermansia

**eTable2**

| | |
|---|---|
| 1 | k__Bacteria;p__[Thermi];c__Deinococci;o__Thermales;f__Thermaceae;g__Thermus |
| 2 | k__Bacteria;p__Proteobacteria;c__Alphaproteobacteria;o__Rhizobiales;f__Methylobacteriaceae;g__Methylobacterium |
| 3 | k__Bacteria;p__[Thermi];c__Deinococci;o__Deinococcales;f__Deinococcaceae;g__Deinococcus |
| 4 | k__Bacteria;p__Proteobacteria;c__Alphaproteobacteria;o__Rhizobiales;__;__ |
| 5 | k__Bacteria;p__Proteobacteria;c__Alphaproteobacteria;o__Rhizobiales;f__Rhizobiaceae;__ |
| 6 | k__Bacteria;p__Proteobacteria;c__Alphaproteobacteria;o__Rhizobiales;f__Rhizobiaceae;g__Agrobacterium |
| 7 | k__Bacteria;p__Proteobacteria;c__Alphaproteobacteria;o__Rhizobiales;f__Bradyrhizobiaceae;__ |
| 8 | k__Bacteria;p__Proteobacteria;c__Alphaproteobacteria;o__Rhizobiales;f__Brucellaceae;g__Ochrobactrum |
| 9 | k__Bacteria;p__Proteobacteria;c__Alphaproteobacteria;o__Rhizobiales;f__Methylobacteriaceae;__ |
| 10 | k__Bacteria;p__Firmicutes;c__Bacilli;o__Bacillales;f__[Thermicanaceae];g__Thermicanus |
| 11 | k__Bacteria;p__Proteobacteria;c__Alphaproteobacteria;o__Rhizobiales;f__Hyphomicrobiaceae;g__Devosia |
| 12 | k__Bacteria;p__Proteobacteria;c__Alphaproteobacteria;o__Rhizobiales;f__Beijerinckiaceae;g__Chelatococcus |
| 13 | k__Bacteria;p__Proteobacteria;c__Alphaproteobacteria;o__Rhizobiales;f__Aurantimonadaceae;__ |
| 14 | k__Bacteria;p__Proteobacteria;c__Alphaproteobacteria;o__Rhizobiales;f__Bartonellaceae;g__ |
| 15 | k__Bacteria;p__Proteobacteria;c__Alphaproteobacteria;o__Rhizobiales;f__Hyphomicrobiaceae;__ |
| 16 | k__Bacteria;p__Proteobacteria;c__Alphaproteobacteria;o__Rhizobiales;f__Methylobacteriaceae;g__ |
| 17 | k__Bacteria;p__Proteobacteria;c__Alphaproteobacteria;o__Rhizobiales;f__Phyllobacteriaceae;g__Mesorhizobium |
| 18 | k__Bacteria;p__[Thermi];c__Deinococci;o__Deinococcales;f__Trueperaceae;g__Truepera |
| 19 | k__Bacteria;p__WPS-2;c__;o__;f__;g__ |
| 20 | k__Bacteria;p__Proteobacteria;c__Alphaproteobacteria;o__Rhizobiales;f__Xanthobacteraceae;__ |
| 21 | k__Bacteria;p__BRC1;c__PRR-11;o__;f__;g__ |
| 22 | k__Bacteria;p__Proteobacteria;c__Alphaproteobacteria;o__Rhizobiales;f__Hyphomicrobiaceae;g__Rhodoplanes |
| 23 | k__Bacteria;p__Proteobacteria;c__Alphaproteobacteria;o__Rhizobiales;f__Phyllobacteriaceae;__ |
| 24 | k__Bacteria;p__Proteobacteria;c__Alphaproteobacteria;o__Rhizobiales;f__Bradyrhizobiaceae;g__Bosea |
| 25 | k__Bacteria;p__Proteobacteria;c__Alphaproteobacteria;o__Rhizobiales;f__Hyphomicrobiaceae;g__Pedomicrobium |
| 26 | k__Bacteria;p__Proteobacteria;c__Alphaproteobacteria;o__Rhizobiales;f__Phyllobacteriaceae;g__Aliihoeflea |
| 27 | k__Bacteria;p__Proteobacteria;c__Alphaproteobacteria;o__Rhizobiales;f__Rhizobiaceae;g__Rhizobium |

**eTable3**

| Patient Type 1 | Patient Type 2 | Shannon Entropy U-Test | Bray-Curtis PERMANOVA |
|---|---|---|---|
| BPD | Inpatient | 0.068 | 0.003 |
| BPD | Outpatient Older | 0.212 | 0.022 |
| BPD | Outpatient Younger | 0.051 | 0.003 |
| Inpatient | Outpatient Older | 0.399 | 0.145 |
| Inpatient | Outpatient Younger | 0.795 | 0.191 |
| Outpatient Older | Outpatient Younger | 0.381 | 0.123 |
